# Functional imaging and quantification of multi-neuronal olfactory responses in *C. elegans*

**DOI:** 10.1101/2022.05.27.493772

**Authors:** Albert Lin, Shanshan Qin, Helena Casademunt, Min Wu, Wesley Hung, Greg Cain, Nicolas Z. Tan, Raymond Valenzuela, Leila Lesanpezeshki, Vivek Venkatachalam, Cengiz Pehlevan, Mei Zhen, Aravinthan D.T. Samuel

**Affiliations:** Department of Physics, Harvard University, Cambridge, MA, USA; John A. Paulson School of Engineering and Applied Sciences, Harvard University, Cambridge, MA, USA; Center for Brain Science, Harvard University, Cambridge, MA, USA; Lunenfeld-Tanenbaum Research Institute, Mount Sinai Hospital, Toronto, ON, Canada; Department of Physics, Northeastern University, Boston, MA, USA

**Keywords:** C. elegans, multi-neuronal imaging, chemosensation, odor representation

## Abstract

Many animals perceive odorant molecules by collecting information from ensembles of olfactory neurons. Each neuron employs receptors that are tuned to recognize certain odorant molecules by chemical binding affinity. Olfactory systems are able, in principle, to detect and discriminate diverse odorants by using combinatorial coding strategies. Multineuronal imaging with high-throughput stimulus delivery allows comprehensive measurement of ensemble-level sensory representations. We have used microfluidics and multineuronal imaging to study ensemble-level olfactory representations at the sensory periphery of the nematode *C. elegans*. The collective activity of nematode chemosensory neurons reveals high-dimensional representations of olfactory information across a broad space of odorant molecules. We reveal diverse tuning properties and dose-response curves across chemosensory neurons and across odorants. We describe the unique contribution of each sensory neuron to an ensemble-level code for volatile odorants. We also show how natural stimuli, a set of nematode pheromones, are encoded by the sensory periphery. The integrated activity of the *C. elegans* chemosensory neurons contains sufficient information to robustly encode the intensity and identity of diverse chemical stimuli.

## Introduction

Many animals exhibit diverse behaviors – navigating the world, finding food, avoiding dangers – on the basis of diverse olfactory cues. To do this, their olfactory systems must distinguish the identity and intensity of numerous odorant molecules.

Insect and mammalian olfactory systems employ large ensembles of olfactory sensory neurons to detect and distinguish volatile odorants and pheromones (1–6). Each olfactory sensory neuron usually expresses a specific olfactory receptor that confers the neuron’s sensitivity to volatile odorant molecules. Each receptor is tuned to recognize odorant molecules by chemical binding affinity (7). A given receptor is typically activated by many different odorant molecules, each to varying degree reflecting differences in chemical affinity. A given odorant molecule also typically activates multiple olfactory receptors to varying degrees (1,8). This allows olfactory systems as a whole to detect and discriminate large varieties of odorant molecules, suggestive of combinatorial coding strategies.

Olfaction is an essential sensory modality in *C. elegans*. The nematode is sensitive to many odorants across a wide range of concentrations (9–11). Compared to larger animals, nematode olfactory circuits have a compact and distinct cellular and molecular organization. The *C. elegans* genome encodes >1000 putative chemosensory GPCR receptors, suggesting a substantial capacity for olfactory detection (12,13). This large receptor family is expressed in a small nervous system with only 11 pairs of amphid chemosensory neurons (12,14). These chemosensory neurons are often characterized as sensors for specific modalities including the detection of volatile odorants (AWA, AWB, AWC) (15–21), soluble chemicals (ASE) (22,23), as-caroside pheromones (ADL, ASK, ADF) (24–29), and nociception (ASH) (10,30–33). Some chemosensory neurons are also polymodal, detecting gases (CO_2_, O_2_) or temperature changes in addition to volatile and soluble odorants (12,34).

The neurons AWA, AWB, and AWC are thought to be the primary detectors of volatile odorants. Laser ablation of AWA or AWC significantly degrades chemotaxis towards selected attractive volatile odorants (35). However, chemotaxis is not completely abolished. Even when both AWA and AWC are ablated, animals are still able to move towards odorant sources. Similar experiments with selected organic compounds and salts showed that ablation of other chemosensory neurons — ASE, ADF, ASG, ASI, ASJ, and ASK — degrades chemotaxis to a lesser extent (10). Thus, the loss of certain neurons impacts the behavioral response to certain odorants more severely. These early results suggest that – although some neurons are more important for chemotaxis towards certain odorants than others – C. *elegans* chemosensation does not rely on signals from single neurons.

We now have a rich understanding of the stimulus-evoked properties of chemosensory neurons in *C. elegans*. Most studies have probed the detection of selected odorant molecules by individual chemosensory neurons (15–21). Isoamyl alcohol is detected by AWC, AWB, and ASH (15). Diacetyl is detected by AWA at low concentrations and by ASH at high concentrations, with AWA also responding to a wide range of volatile odorants (16). Benzaldehyde is detected by AWA, AWB, AWC, and ASE (17). In some cases, the left and right pairs of a chemosensory neuron type detect different odorant molecules. A library of single-neuron labeled lines has been used to assess single-neuron responses to a multimodal panel of stimuli, including two volatile odorants, isoamyl alcohol and diacetyl, at one concentration (18). This study reported sparse activation of chemosensory neurons.

The most thoroughly characterized odorant receptor in *C. elegans* is ODR-10, expressed in AWA and shown to respond to diacetyl (an attractive stimulus). Ectopic expression of ODR-10 in AWB leads to diacetyl repulsion, suggesting that AWA and AWB may be linked to attractive and aversive behaviors, respectively (36). AWB and AWC have also been found to be necessary for aversive olfactory learning (37). AWA neurons fire action potentials that may encode stimulus-specific features (38). Complex activity patterns of single neurons such as AWA have been directly mapped to behavioral patterns (16,17,36,38,39).

The left and right AWC neurons, AWCL and AWCR, are stochastically asymmetric. In each worm, one neuron (either AWCL or AWCR) adopts the identity AWC^ON^ and its lateral pair adopts the identity AWC^OFF^ (20). AWC^ON^ detects butanone, while AWC^OFF^ detects 2,3-pentanedione (40,41). Like AWA, AWC has been shown to have complex single-neuron properties, capable of changing its response properties in a context-dependent manner (42,43). ASE neurons, primarily characterized as gustatory neurons, also respond asymmetrically to different ions during salt chemotaxis. ASEL detects sodium ions and ASER detects chloride and potassium ions (22,23). It is not known whether ASE has any asymmetric responses to volatile odorants.

Sensory adaptation has been observed in AWC, ASH, and ASE. When presented with a prolonged chemical stimulus from minutes to hours, neuronal activity is gradually reduced (16,44–47).

Although odorant-evoked responses in many chemosensory neurons in *C. elegans* have been well characterized, how their collective dynamics might represent odorant information as an ensemble remains unexamined. We set out to characterize how the chemosensory ensemble in *C. elegans* encodes a chemically diverse space of volatile odorants at different concentrations, and to understand the tuning properties of each chemosensory neuron with respect to this large odorant space and ensemble-level code.

We assembled a panel of olfactory stimuli spanning a diverse molecular chemistry and used microfluidics to deliver these odorants at multiple concentrations (**Figure 1B**). To efficiently record neuronal responses at the sensory periphery, we used a transgenic animal that allowed the simultaneous measurement of intracellular calcium dynamics in all amphid chemosensory neurons (**Figure 1C**).

**Figure 1.**
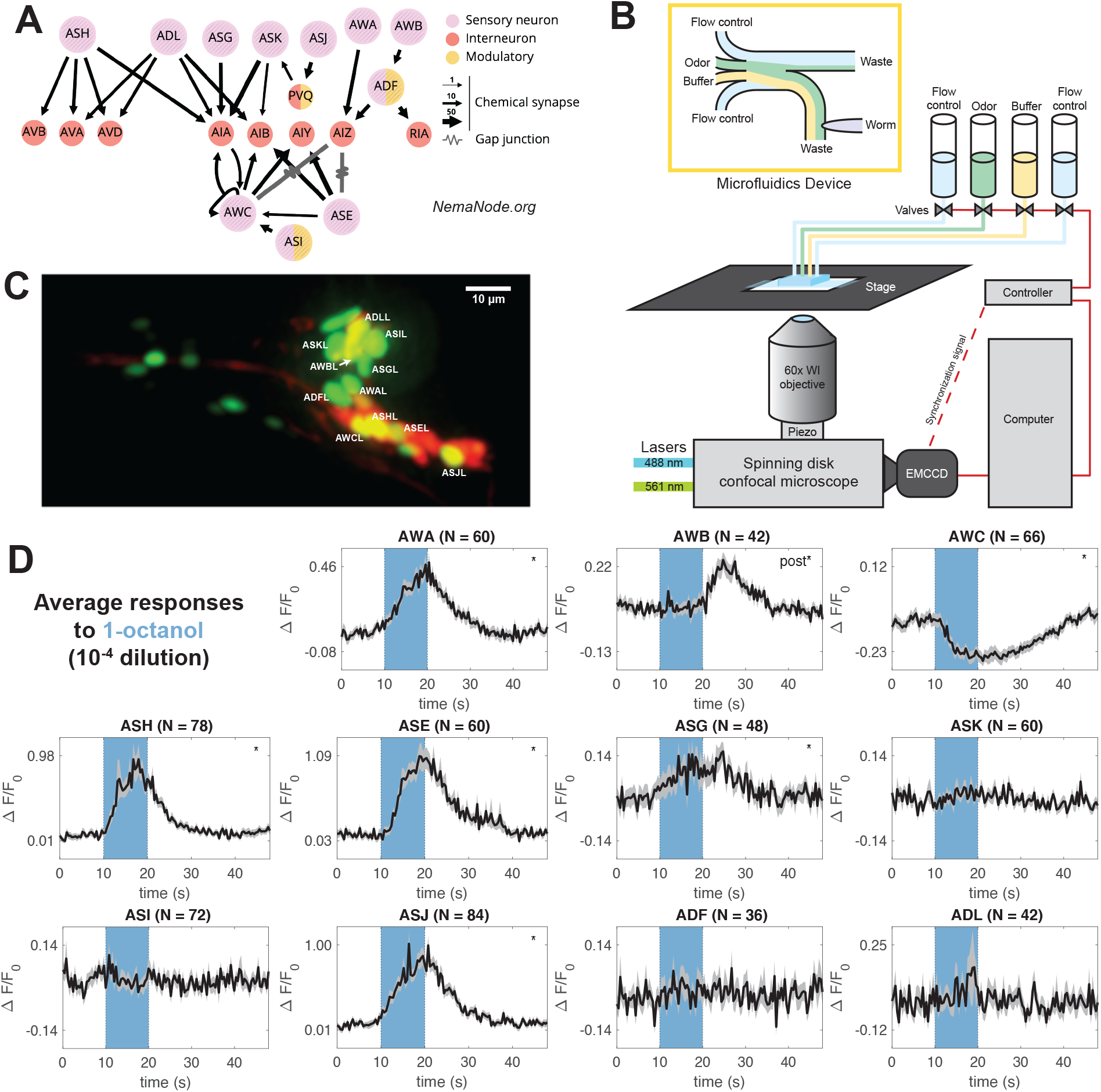
Labeling and recording from chemosensory neurons. **(A)** Downstream partners of the 11 chemosensory neurons in the *C. elegans* connectome (48,49). Panel generated at nemanode.org. **(B)** Adult *C. elegans* were immobilized inside a microfluidic device and controllably presented with odorant solutions. Each animal was volumetrically imaged at 2.5 Hz with a spinning disk confocal microscope during stimulus presentations. **(C)** Animals expressed nuclear-localized GCaMP6s in all ciliated sensory neurons. A sparse wCherry landmark distinguished the 11 chemosensory neurons. Here, a dual-color maximum projection image shows the head of the worm. The 11 chemosensory neurons on the near (L) side are labeled. For clarity, the chemosensory neurons on the far side and other ciliated neurons are not labeled. **(D)** Neuronal activity traces of the 11 chemosensory neurons in response to a single odorant presentation (1-octanol, 10^−4^ dilution), averaged across trials. The 10s odorant delivery period is shown by the colored bar. Significant responses (*q* ≤0.01) are marked with stars, with “post” indicating a significant response to stimulus removal (OFF response). Error bars (gray) are standard error of the mean.

We found that most odorant-evoked responses are widespread across the chemosensory ensemble. Dose-response curves are different for different odorant molecules, whether comparing the responses of the same neuron to different odorants or comparing the responses of different neurons to the same odorant. Odorant identity and intensity information can be reliably decoded by the collective activity of the chemosensory ensemble. A set of pheromones also evokes ensemble-level responses, but with a distinct pattern from volatile odorants.

The small nervous system of *C. elegans* has the capacity to use ensemble-level representations to robustly discriminate the identity and intensity of odorant molecules across olfactory stimulus space.

## Results

### Calcium imaging of chemosensory neurons with representative odorant stimuli

We developed a GCaMP6s calcium reporter line to simultaneously record calcium dynamics in all ciliated sensory neurons (**Methods**). In this study, we focus on the 11 pairs of amphid chemosensory neurons: AWA, AWB, AWC, ASE, ASG, ASH, ASI, ASJ, ASK, ADL, and ADF (**Figure 1A**). We immobilized and positioned young adult *C. elegans* in a microfluidic device that allows odorants to flow past its nose (**Figure 1B**) (50). We adapted a multichannel microfluidic device (4) to control the delivery of pulses of single and mixed odorant solutions. Volumetric imaging was performed at 2.5Hz with a spinning disk confocal microscope (**Figure 1D**).

We assembled a stimulus panel of 23 odorant molecules, selected from 122 molecules that had been previously used to study *C. elegans* olfaction (35,51). The 23 odorant molecules were chosen to span the chemical diversity of previously used stimuli. The panel includes exemplars of six chemical classes: alcohols (1-pentanol, 1-hexanol, 1-heptanol, 1-octanol, 1-nonanol, isoamyl alcohol, and geraniol), aromatics (benzaldehyde and methyl salicylate), esters (ethyl acetate, ethyl butyrate, pentyl aetate, ethyl heptanoate, and butyl butyrate), ketones (2-butanone, diacetyl, 2-heptanone, 2-nonanone, and 2,3-pentanedione), pyrazines (2,5-dimethyl pyrazine and 2-methyl pyrazine), and thiazoles (2-isobutylthiazole and 2,4,5-trimethylthiazole). To assess chemical diversity, we constructed a geometrical odor space on the basis of physical and chemical descriptors of molecular structure (52). Our 23 odorants broadly sample this geometrical space (**Figure S3A**) (52).

We recorded the responses of all amphid chemosensory neurons to >70 stimulus conditions, testing each of the 23 odorants at multiple concentrations. Individual animals were repeatedly presented with series of 10s odorant pulses separated by 30s buffer blanks. For each stimulus condition, we recorded the responses to approximately 100 odor presentations across multiple animals (**Figure 2A-C, S3C-D**). The highest concentrations we tested were 10^−4^ dilutions. The lowest concentrations we tested – 10^−8^ dilutions – did not elicit significant responses from any sensory neuron.

**Figure 2.**
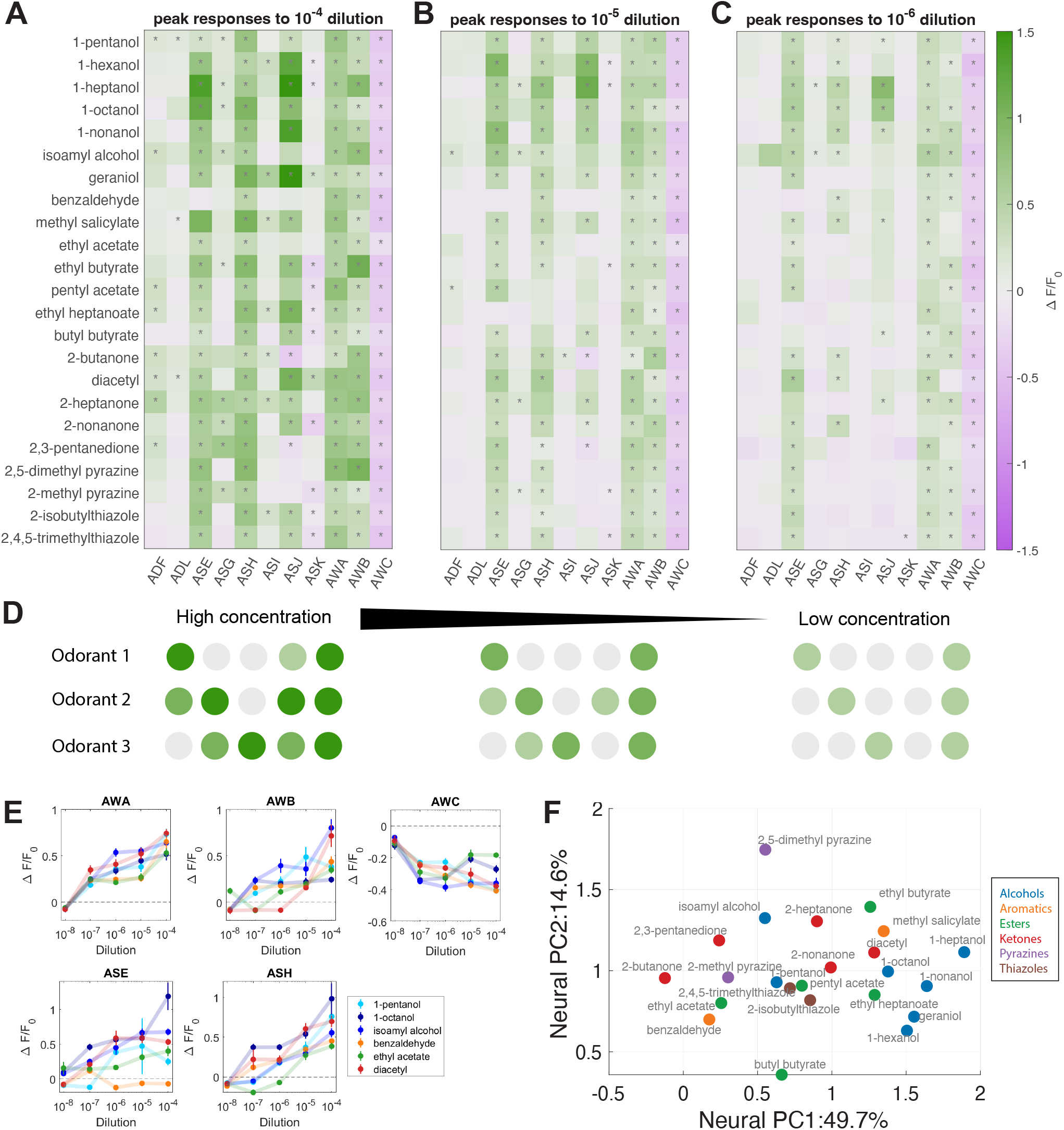
Ensemble responses to a broad odorant panel. Average peak responses of the 11 chemosensory neurons to odorants at **(A)**high concentration (10^−4^ dilution), **(B)** medium concentration (10^−5^ dilution), and **(C)** low concentration (10^−6^ dilution). Peaks were computed from a time window from onset of odor delivery to 10 s after odor removal. Responses are reported as Δ*F*/*F*_0_, and significant responses (*q* ≤ 0.01, 2-tailed, paired t-tests) are indicated with stars. Most odorants elicit significant responses from unique combinations of neurons. **(D)** Schematic of coding strategy observed in panels **A-C**. Different odorants evoke responses in distinct subsets of sensory neurons. Responses are generally stronger at high concentrations. Additional neurons are activated as concentration increases. **(E)** Dose responses of the peak responses of AWA, AWB, AWC, ASE, and ASH are diverse, with distinct concentration-dependent curves in response to different odorants. See **Figure S3F** for dose responses of the other 6 sensory neurons. Error bars are standard error of the mean. **(F)** A PC space built from standardized peak average neural responses. Chemical class is indicated by color. Some odorant classes, such as alcohols and ketones, have more similar neural representations, while other odorant classes, such as esters, have more diverse representations. Refer to **Figure S3H** for PC loadings.

### Odorants elicit ensemble responses

Across our odorant panel, calcium imaging captured many sensory neuron responses, some previously characterized and some unknown. Nearly every odorant reliably activated more sensory neurons than previously described. For example, the odorant diacetyl, attractive at low concentrations, reliably activated AWA upon odor onset (53)) at all concentrations (**Figure 2A-C**). The odorant 1-octanol, a repellent, reliably activated ASH and inhibited AWC (54) across concentrations. We discovered that additional sensory neurons also reliably responded to both odorants. For example, AWC was inhibited by diacetyl and ASJ was activated by 1-octanol. Isoamyl alcohol not only activated AWA, AWB, AWC and ASH at different concentrations, as previously reported (15), but also activated the ASE and ASG neurons (**Figure 2A-C**). At high concentrations, every odorant elicited responses from multiple sensory neurons. We observed significant overlap in the sets of responding neurons for different odorants (**Figure 2A-D**).

Most chemosensory neurons exhibited excitatory responses – increases in intracellular calcium levels during stimulus presentation. Some neurons exhibited inhibitory responses – decreases in intracellular calcium levels below the baseline level. Previous work has shown that AWC is inhibited by several odorants in our panel, including diacetyl, benzaldehyde, and 2-butanone (16,17,21,37). In our stimulus conditions, AWC is inhibited by every odorant in our panel (**Figure 2A**). We also discovered that ASK is inhibited by many odorants including ethyl butyrate and 2-nonanone (**Figure S2B**). Some neurons are inhibited by certain odorants but excited by others. For example, ASJ is strongly inhibited by 2-butanone but strong excited by 1-nonanol (**Figure S2C**).

Most chemosensory neurons exhibited ON responses to most odorants – changes in calcium levels upon odorant onset. We also observed OFF responses – changes in calcium levels upon odorant removal. For example, AWB has been reported to exhibit ON and OFF responses at different isoamyl alcohol concentrations (15). We confirmed this result, and also found that AWB had ON responses to some odorants, such diacetyl at high concentration, and OFF responses to others, such as 1-hexanol and 1-octanol (**Figure 1D, S2A**).

The left and right ASE neurons exhibited strong asymmetry in their responses to two odorants in the panel: heptanoate and butyl butyrate both activated ASEL and inactivated ASER (**Figure S2D**). The ASE neurons were previously shown to respond asymmetrically to non-volatile chemical stimuli (20,23). AWC, another pair of neurons with known structural asymmetry (20), might exhibit moderate differences in their response dynamics when presented with short odorant pulses (21). Whereas the cellular identities of ASEL and ASER are defined by their handedness, AWCL and AWCR stochastically adopt the identities of AWC^ON^ or AWC^OFF^ (AWC^ON^ can be identified via cell-specific expression of the *str-2* promoter). Here, we cannot distinguish which neuron is AWC^ON^ or AWC^OFF^, except by inference from neuronal activity patterns. Because all other left and right sensory neurons respond symmetrically to all odorants, and because the left and right ASE and AWC neurons also respond symmetrically to many odorants, we grouped signals from left and right sensory neurons in all analyses unless otherwise noted.

To compare the temporal dynamics of chemosensory neurons across odorants, we computed pair-wise cross-correlations of the activity time courses for each odorant (**Figure S4A-B**). We found that matrices of pairwise cross-correlations are distinct for different odorants. From both peak responses and dynamics, the diversity of ensemble-level dynamics is as large as the number of tested odorants. The compact sensory neuron ensemble of *C. elegans* may be able to encode the identities of numerous odorants by using the combinatorially large space of distinct activity patterns.

### Sensory representations are not dependent on synaptic connections

The *C. elegans* connectomes have revealed consistent axoaxonic chemical synapses between some sensory neurons and from some interneurons to sensory neurons (**Figure 1A**) (48,49). These connections raise the possibility that ensemble representations might not entirely reflect independent responses from individual neurons.

We examined this possibility by analyzing ensemble responses in an *unc-13(s69)* mutant where synaptic vesicle fusion is nearly fully blocked (55) (**Figure 3A**). We sampled five odorants that represent different chemical classes. In all cases, nearly identical groups of neurons significantly responded (*q* ≤ 0.01) in wild-type and *unc-13* mutants (**Figure 3B**).

**Figure 3.**
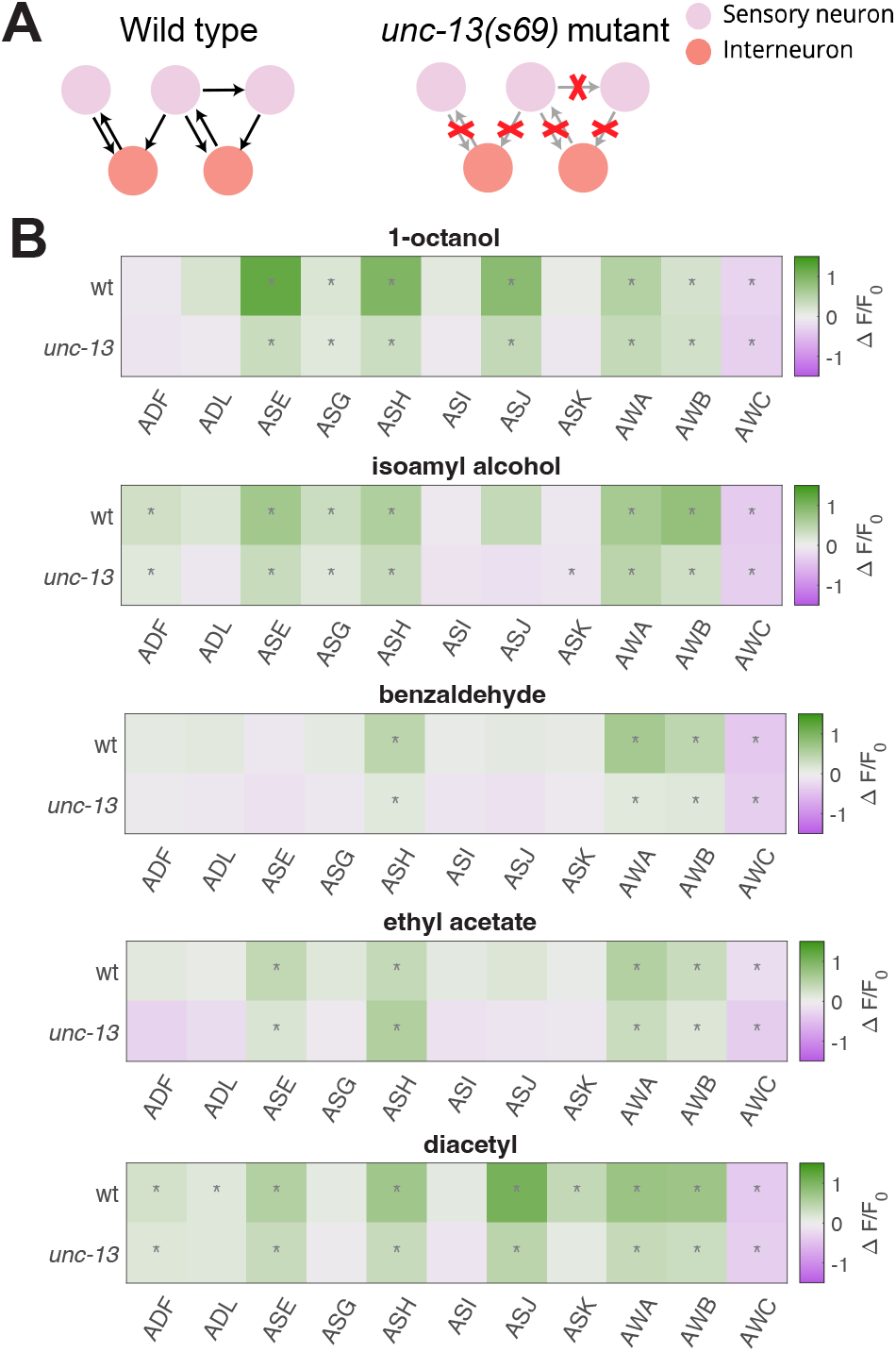
Odorant representations in synaptic transmission mutants. **(A)** The majority of chemical synapses in *unc-13(s69)* synaptic transmission mutants are non-functional. We recorded neural activity in these mutants during odor presentation. **(B)** When presented with the same odorants, similar sets of neurons significantly (*q* ≤ 0.01) responded in wild-type and *unc-13* mutants.

Chemical synaptic transmission does not appear to be the dominant factor in ensemble responses – similar neuronal ensembles respond to diverse olfactory stimuli in animals with or without chemical synaptic communication. The tuning of each neuron to an odorant is likely to be cell intrinsic, a function of the receptors expressed in each neuron.

### Olfactory representations broaden with increasing concentrations

To compare the response properties of different neurons, we constructed dose-response curves for all 11 chemosensory neurons in response to odorants from our panel over 3-5 orders of magnitude in concentration (**Figure 2A-C, S3C-D**).

For most odorants and neurons, response magnitudes increased monotonically with odorant concentration – neurons activated at low concentrations were also activated at all higher concentrations. Every odorant is associated with a characteristic set of neurons activated at all concentrations above the detection threshold. Across all concentrations, for example, 1-pentanol activates AWA and AWC; 1-octanol activates ASE, ASH, AWA, AWB and AWC; and benzaldehyde activates AWA, AWB and AWC. Each set of responding neurons may constitute a unique olfactory representation associated with each odorant identity.

For many odorants, increasing concentration spatially broadens olfactory representation by activating more sensory neurons. Different neurons exhibit different thresholds for different odorants. For example, AWB is only activated by 1-pentanol at concentrations above 10^−5^ dilution, and ADF, ADL, and ASG are only significantly activated by 1-pentanol at 10^−4^ dilution, the highest tested concentration (**Figure S3E**). Thus, odorant intensity is represented partly by the magnitude of responses of activated neurons and partly by the number and identities of activated neurons (**Figure 2D**).

We used phase-trajectory analysis to illustrate the temporal dynamics of ensemble-level odorant representations. In a low-dimensional principal component space, these representations follow closed trajectories as they evolve over time following odor presentation (**Figure S4C**). Along each trajectory, neurons become activated, reach their peak responses, and return to baseline. In this space, the responses to different odorants follow trajectories with different headings from the origin. Trajectories for responses to the same odorant at different concentrations are aligned in direction but differ in magnitude.

### Diversity in dose responses across neurons and odorants

The dose-response curves of the 11 chemosensory neurons exhibit significant diversity (**Figure 2E, S3F**). Each odorant can evoke dose-response curves with different steepnesses and thresholds in different neurons. Conversely, each sensory neuron can exhibit dose-response curves with different steepnesses and thresholds for different odorants.

In some cases, neurons detected an odorant with slowly graded responses over a broad dynamic range. Graded responses include AWA’s response to 1-pentanol and AWB’s response to 1-heptanol (**Figure 2E, S3F**). In other cases, neurons exhibited steep response functions, becoming fully activated or fully inhibited above a sharply defined threshold. Step-like responses include ASE’s response to 1-pentanol and AWB’s response to 1-octanol.

Diversity in dose response curves contrasts with insects and mammals, where olfactory sensory neurons typically exhibit similar dose response curves across neurons and across odorants (4,56,57). In insects and mammals, each sensory neuron is generally equipped with one receptor type, whereas in *C. elegans* each neuron likely expresses multiple receptors (12,13). For a given nematode neuron, the presence of receptors to multiple odorants, each with different kinetics, may explain dose response diversity.

### Comparing ensemble-level representations of chemically similar odorants

Different odorants activate distinct but overlapping subsets of the chemosensory ensemble (**Figure 2A-C**). Quantitative differences in the sensitivity of chemosensory neurons to odorants will depend on cell-specific patterns of receptor expression. In most olfactory systems, a typical olfactory receptor is activated by a range of structurally similar odorant molecules with common chemical features. This leads to a systematic dependence of ensemble-level olfactory representations on odorant chemistry. To assess this dependence in *C. elegans*, we performed hierarchical clustering of odorants from our panel based on ensemble-level responses evoked at high concentrations (**Figure S3G**). The representations of some molecular classes clustered together. For example, ensemble-level responses to a set of straight-chain alcohols (1-hexanol, 1-heptanol, 1-octanol, and 1-nonanol) were similar to one another, and the ensemble-level response to a set of ketones (2-butanone, 2,3-pentanedione, and 2-heptanone) were likewise similar. On the other hand, the esters in our panel, a group more diverse in their chemical structure, produced a broader set of representations.

Principal Components Analysis (PCA) is a quantitative means of assessing the similarity of high-dimensional ensemble-level representations. We constructed a principal component space from all average ensemble-level peak responses, and asked how different odorants are distributed in this space. Consistent with observations from hierarchical clustering, responses to certain classes of odorants, such as alcohols and ketones, are close to each other in PC space. Responses to members of other classes, such as esters, are distributed more broadly (**Figure 2F**). The loading of the first three principal components of this space allows us to assess the relative contribution of each sensory neuron to ensemble representations (**Figure S3H**). We observed a broad distribution of principal component loading, a measure that suggests that all chemosensory neurons contribute to the separability of odorant representations.

### Sensory neurons are broadly or narrowly tuned in chemical space

How are individual sensory neurons tuned in chemical space? Olfactory sensory neurons are tuned to odorants by the relative binding affinities of receptors for different ligands (58). In animals where sensory neurons express single receptor types, this leads to a systematic dependence of ensemble representation on the chemical properties of the odorant and the receptors (4,59–61). In *C. elegans*, the tuning of a sensory neuron may also be shaped by the expression of multiple different receptors. To explore the tuning of the sensory neurons in chemical space, we projected the activity of each neuron into the space of chemical structure based on molecular descriptors of each odorant (**Figure S3A**) (52).

We observed both broad and narrow tuning among sensory neurons. For example, AWA, AWB, AWC, and ASE are broadly tuned, each responding to most tested odorants at high concentrations (**Figure 2A-C, 4A**). In contrast, ADF, ADL, ASG, ASI, ASJ, and ASK are narrowly tuned, each responding to a small set of odorants even at the highest tested concentrations.

**Figure 4.**
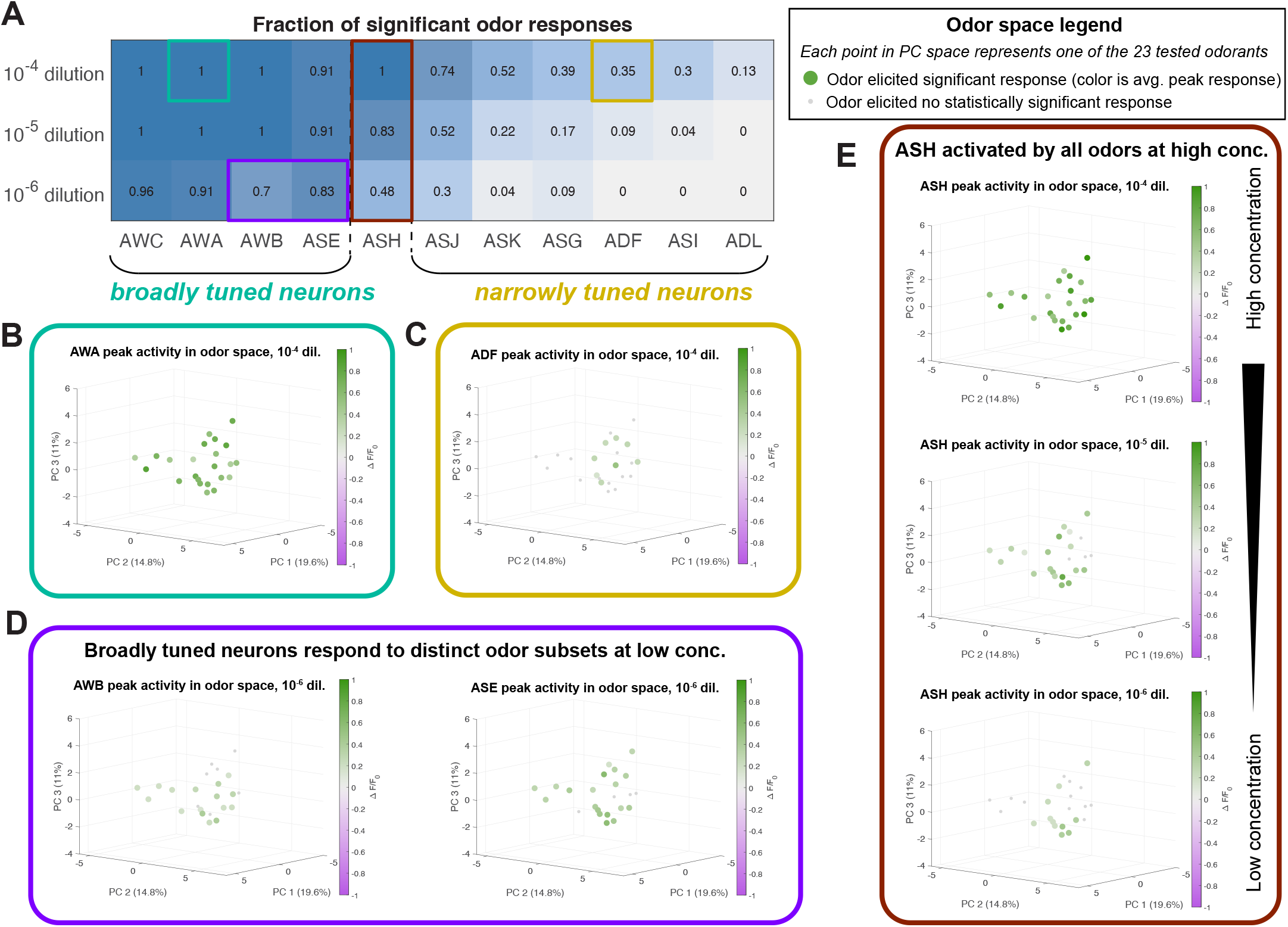
Chemosensory neuron tuning. **(A)** The fraction of odorants in our 23-odor panel which elicited significant responses (*q* ≤ 0.01) in each neuron, at three different concentrations. We consider neurons which responded to the majority of presented odors as “broadly tuned”, and neurons which responded to a small numbers of odors as “narrowly tuned”. For each neuron, we plot peak responses to odorants in a space constructed from chemical descriptors (**Figure S3A**). **(B)** The activity of broadly tuned neurons (ex: AWA) spans this space, while **(C)** the activity of narrowly tuned neurons (ex: ADF) is confined to a subset of chemically similar odorants. **(D)** At low concentrations, broadly tuned neurons respond to distinct subsets of odorants. **(E)** ASH, a polymodal nociceptor, is activated by all tested odorants at high concentration, but is only activated by a small set of repulsive odorants at low concentration. See **Figure S5** for these plots for all neurons.

ASH is broadly tuned at high concentrations and narrowly tuned at low concentrations (**Figure 4A**), a pattern that might reflect its role as a nociceptor, mediating avoidance of any odorant when delivered at a sufficiently high concentration (**Figure 2A-C**). In previous behavioral experiments, most odorants in our panel were shown to be attractive at low concentrations and aversive at high concentrations. A few odorants – 1-heptanol, 1-octanol, and 1-nonanol – are aversive at any tested concentration (**Appendix B**). The odorants to which ASH is most sensitive are those that are aversive at all concentrations.

The responses of each sensory neuron occupy contiguous domains in chemical space. Each domain encompasses chemically similar odorant molecules that are effective stimuli for each sensory neuron (**Figures 4B-E, S5**). At high concentrations, broadly tuned neurons – such as AWA – extend responses throughout the chemical structure space. Even at high odorant concentrations, narrowly tuned neurons – such as ADF – extend responses over a smaller contiguous region of chemical space.

At lower concentrations, most broadly tuned neurons extend responses over a smaller region of chemical space, revealing structural characteristics of molecules to which each sensory neuron is most sensitive. AWA is most strongly activated by ketones, AWB is most strongly activated by esters, and ASE is most activated by alcohols (**Figure S5**). At low concentrations, ASH responds to odorants throughout chemical space, a breadth that may reflect the fact that any odorant delivered at sufficiently high concentration is repellent. The observation that each sensory neuron extends its sensitivity range across a contiguous region of chemical structural space suggests that each neuron is tuned to shared molecular properties of a set of odorant stimuli, as opposed to being faithful ‘labeled-line’ detectors of specific odorants.

### Single-trial responses suffice for discriminating odorant pairs

We observed trial-to-trial variability in odorant responses, both across animals and across odor presentations to the same animal. A potential advantage of ensemble-level coding is additional robustness when discriminating odorants.

We compiled all single-trial responses to each odorant across all datasets. In some recordings where data from individual neurons was missing, we imputed missing activity patterns using the rest of the ensemble (**Appendix D**, **Figure S6A-D**). We used two independent dimensionality reduction methods to visualize the space spanned by single-trial responses – PCA and Uniform Manifold Approximation and Projection (UMAP). In a PC space constructed from the peak responses of all single trials, chemically similar odorants exhibit more similar representations (**Figure S6D**) and chemically dissimilar odorants exhibit dissimilar representations (**Figure S6E**). Overlap in a low-dimensional PC space is an imperfect measure for odorant discrimination because <60% of variance is explained by the first three principal components. Plotting all single-trial responses to all 23 odorants in UMAP space, trials for the same odorant also cluster together, although it is difficult to segregate trials for different odorants in this 2D representation (**Figure 5A**). Both PCA and UMAP analyses indicate that ensemble-level responses for the same odorant are similar. Both analyses also indicate that ensemble representations are high-dimensional, as reduction to 2 or 3 dimensions removes a significant fraction of the variance.

**Figure 5.**
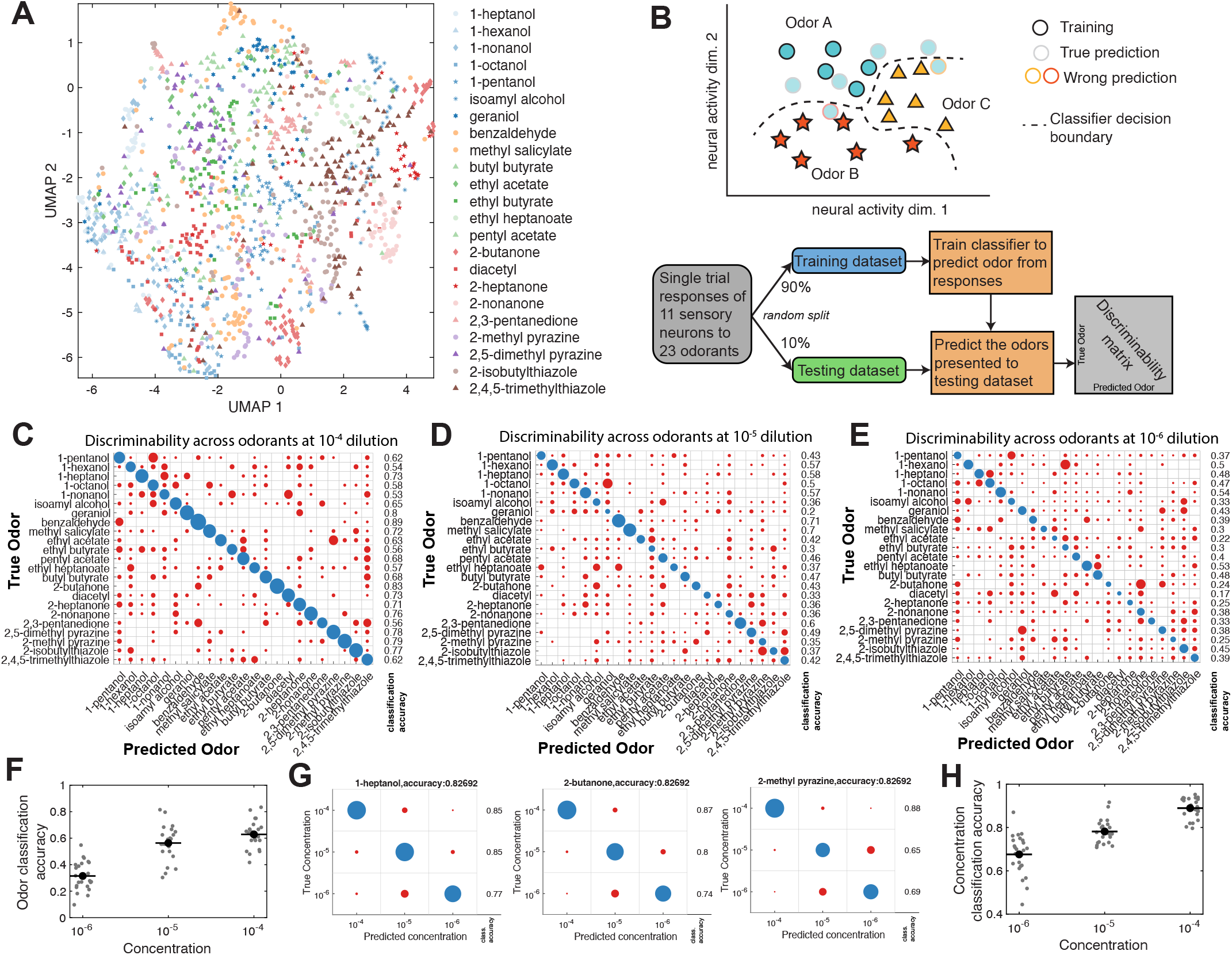
Representative comparisons of single-trial odorant responses. **(A)** A low-dimensional UMAP representation of single-trial neural responses to all 23 odorants at 10^−4^ dilution. Responses to any given odorant generally cluster together. **(B)** Schematic of the multi-class classifier used for theoretical discriminability analysis of single-trial responses. The classifier was trained to predict odor identity from the peak responses of the ensemble of sensory neurons, generating a discriminability matrix. **(C)** Linear discriminability analysis of single-trial peak responses to high-concentration (10^−4^ dilution) odorants, with the presented odorant on the y-axis and the classified odorant on the x-axis. Circle size indicates the number of trials, with correct classifications colored blue and incorrect classifications colored red. The fraction of correctly classified trials for each odorant is to the right. The majority of single trials are correctly classified for each odorant. At lower concentrations, 10^−5^ dilution **(D)** and 10^−6^ dilution **(E)**, classification accuracy diminishes. This is summarized in **(F)**, a scatterplot of multi-class classification accuracy at different concentrations **(C-E)**. **(G)** Within a given odorant (three examples shown), the concentration of the given odorant can be correctly classified based on individual peak responses. **(H)**Across all odorants, concentration classification accuracy at different concentrations is shown.

We asked whether olfactory representations were sufficiently dissimilar for reliable odorant discrimination based on single odorant presentations. To estimate the theoretical discriminability of odorant pairs, we computed errors in binary classification based on the pooled single-trial responses of each odorant pair using logistic regression (**Figure S6F**) and a Support Vector Machine (SVM) (**Figure S6G**). In all cases, binary classification succeeded with low error rate. Thus, any two odorants in our panel are linearly separable based on single-trial ensemble responses.

### Odorant identification based on single-trial responses

We asked whether odorant identity could be uniquely decoded on the basis of single-trial ensemble responses, a task significantly more challenging than binary classification of an odorant pair. We trained a multi-class classifier to perform linear discrimination (**Figure 5B**). We randomly divided all single-trial measurements into a training set (90%) and validation (testing) set (10%). After we trained the classifier with the training set, we tested its performance in predicting odorant identities from single-trial measurements drawn from the validation set (see **Appendix E** for details). This classifier successfully identified odorants in the majority of single-trial measurements at high concentrations (**Figure 5C,F**). Classification accuracy declined at lower odorant concentrations, but succeeded in the plurality of measurements (**Figure 5D-F**).

We used a similar approach to determine whether odorant intensity could be estimated from single-trial measurements. With trained multi-class classifiers, we were able to predict the concentration of a given odorant using single-trial measurements, although accuracy declined at lower concentrations (**Figure 5G-H**). In principle, the ensemble-level spatial map of sensory neuron activity contains sufficient information to determine odorant identity and intensity from single stimulus presentations.

### Virtual neuron knockouts degrade classifier accuracy

To quantify the relative contribution of each sensory neuron to ensemble-level discriminability, we performed virtual knock-outs. We performed virtual knockouts by removing (masking) specific sensory neurons from the dataset and retraining the multi-class classifier on the remaining data. Removing any single sensory neuron led to small decreases in classification accuracy compared to wild-type (**Figure 6B-D**). Classification accuracy was degraded more severely when masking narrowly tuned neurons (such as ASI, ASK, ASJ, and ASG) than masking broadly tuned neurons (such as AWA, ASH, and AWC).

**Figure 6.**
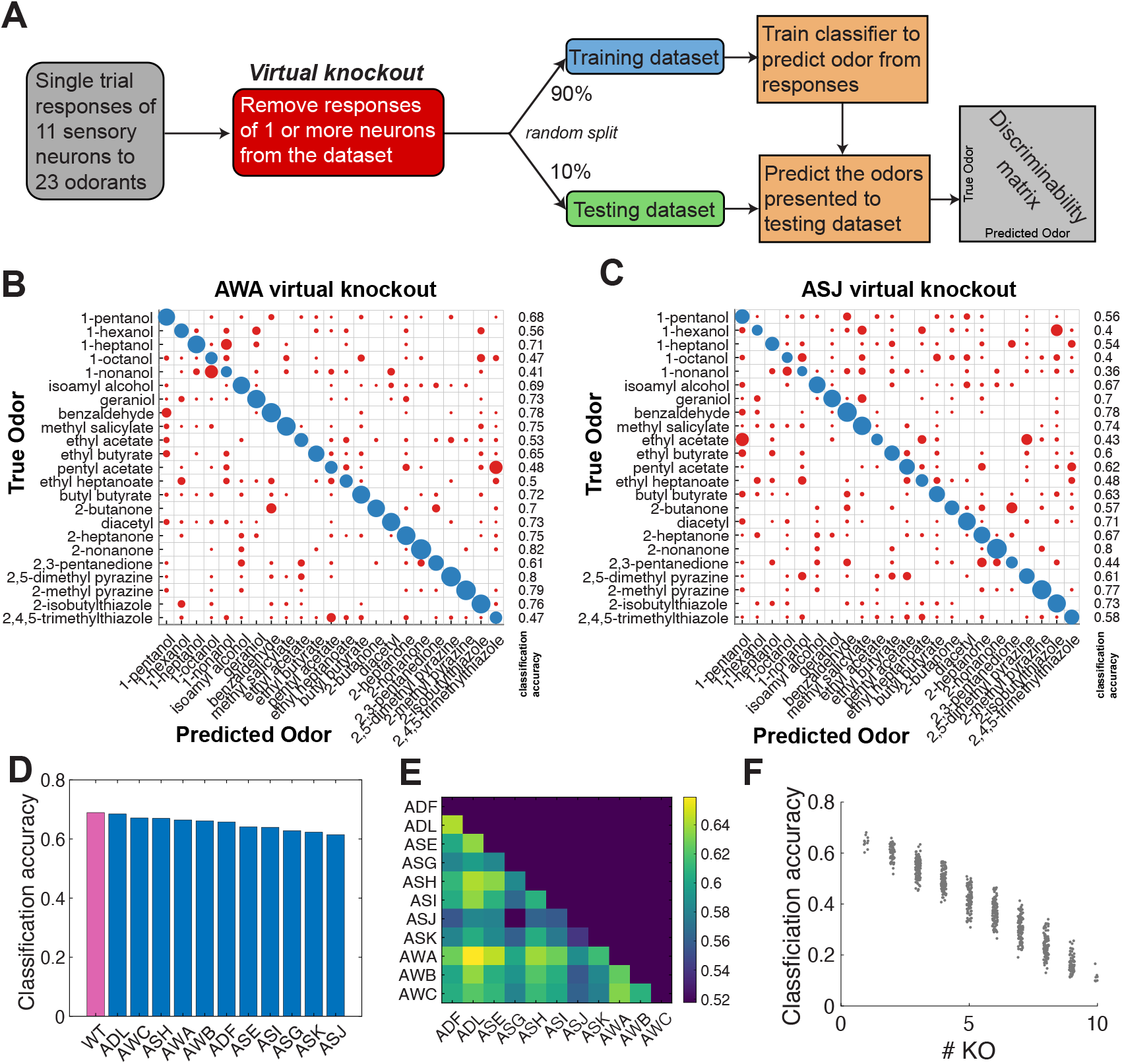
Odorant discriminability is robust to virtual knockouts. **(A)** By removing the responses of 1 or more neurons from the dataset fed into the multi-class classifier, we assess the relative importance of different neurons on the theoretical discriminability of single-trial neural responses. Linear discriminability analysis of single-trial data, with **(B)** AWA or **(C)** ASJ virtually removed from the dataset. Removing different neurons changes the discriminability matrix in different ways. **(D)** We virtually removed each neuron from the dataset, and computed the average classification accuracy for each virtual knockout. Classification accuracy remains close to wild type (all 11 neurons), but is degraded more severely by removal of narrowly tuned neurons (ASI, ASK, ASJ, ASG) than by removal of broadly tuned neurons. **(E)** Virtually removing pairs of neurons produces further reductions in average classification accuracy. **(F)** Plotting average classification accuracy of different sets of virtual knockouts reveals a linear relationship between theoretical classification accuracy and the number of chemosensory neurons.

Masking different neurons degrades the classification accuracy of a given odorant to different degrees. For instance, pentyl acetate is correctly classified 68% of the time when all 11 chemosensory neurons are included. ASJ masking reduces this accuracy to 62%, but AWA masking reduces accuracy to 48%.

Masking any two neurons further decreases average classification accuracy (**Figure 6E**). We computed the average classification accuracy when randomly removing different combinations of multiple neurons. We observed an inverse linear relationship between the number of masked neurons and classification accuracy (**Figure 6F**). Odor identity across olfactory space is thus encoded in a distributed manner across all 11 chemosensory neurons.

### Responses to pheromone stimuli are distinct from those of volatile odorants

All amphid chemosensory neurons are involved in the detection of volatile odorants. We asked whether ensemble-level responses extend to other stimuli. *C. elegans* communicate with each other using pheromones, a mixed group of glycolipid molecules called ascarosides (24,29). We presented youngadult hermaphrodites with a panel of five single ascarosides (#1, #2, #3, #5, and #8) (25).

Similarly to volatile odorants, ascarosides activated multiple sensory neurons (**Figure 7A**). Some neurons – known to respond to ascarosides but narrowly tuned to volatile odorant panel, such as ADL, ADF, and ASK – were strongly activated across our 5 pheromone panel (**Figure 7B**). Pheromones also evoked some activity in neurons that are broadly tuned to volatile odorants. For example, AWA was activated less often by the pheromone panel than by the odorant panel. Thus, pheromone detection may also involve an ensemble-level code, but a code that relies more heavily on those neurons that are narrowly tuned to volatile odorants.

**Figure 7.**
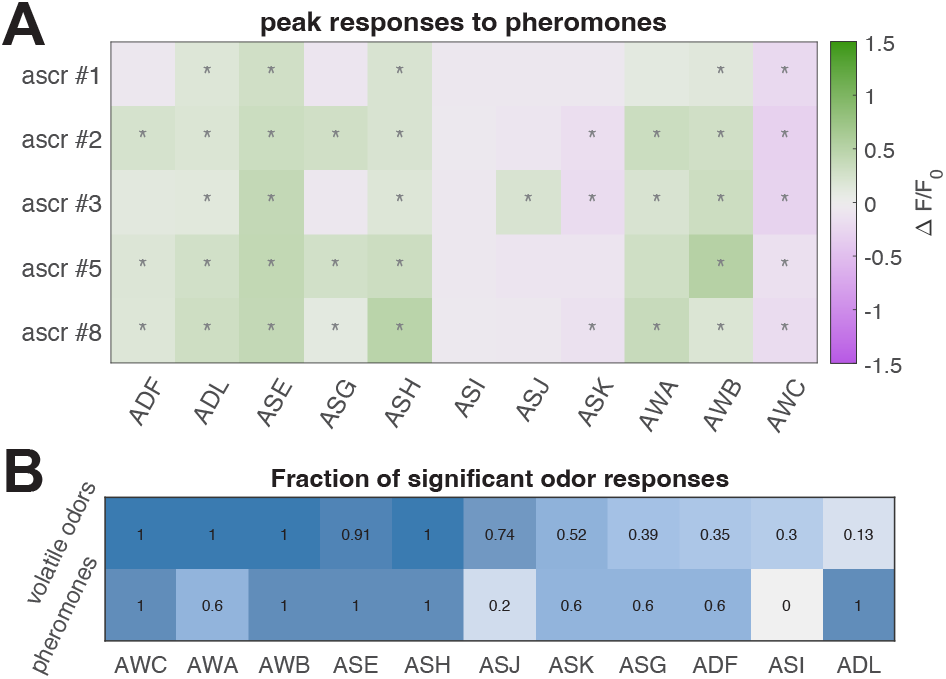
Odorant representations in synaptic transmission mutants and representations of pheromones. **(A)** Average peak responses of the 11 chemosensory neurons to ascaroside pheromones #1, #2, #3, #5, and #8 at a concentration of 200 nM. Responses are reported as Δ*F*/*F*_0_, and significant responses (*q* ≤ 0.01) are indicated with stars. **(B)** The fraction of volatile odorants (out of 23 odorants total) which elicited significant responses in each neuron at high concentration (first row), compared with the fraction of pheromone stimuli (out of 6 stimuli total) which elicited significant responses (second row). Many neurons (such as ADF and ADL) which are narrowly tuned with respect to volatile odorants appear to be activated more often by the ascaroside pheromones.

## Discussion

In insects and vertebrates, the integrated activity of large ensembles of chemosensory neurons is often presumed to enhance odorant discrimination and broaden the space of olfactory perceptions (1–6). The *C. elegans* olfactory system contains only 11 pairs of chemosensory neurons. Each nematode chemosensory neuron is considered a unique class distinguished by dendrite morphologies, wiring partners, and sensory modalities (12,34). Does *C. elegans* integrate information from multiple chemosensory neurons to help discriminate the many different olfactory cues that drive diverse behavioral responses?

We have simultaneously recorded calcium dynamics in all chemosensory neurons in nematodes exposed to a chemically diverse odorant panel. Nearly every distinct odorant stimulus evoked a distinct ensemble-level activity pattern among chemosensory neurons. We show that these highly reproducible ensemble-level patterns can robustly encode odorant identity and intensity throughout a large chemical space.

### *C. elegans* can use its chemosensory neuron ensemble to identify odorants

Previous studies of the *C. elegans* olfactory system largely dissected the properties of individual chemosensory neurons in response to selected odorants (15–21). Many studies have implicitly explored “labeled lines,” where the activity patterns of single sensory neurons are directly mapped to behavioral patterns. Indeed, single olfactory sensory neurons can exhibit complex temporal activity patterns in response to odorant stimulation (16,17,36,38,39,42,43). However, experiments where selected odorants are used to stimulate sensory neurons do not explore how the animal encodes or discriminates diverse olfactory inputs.

We found that most olfactory stimuli activate multiple chemosensory neurons in *C. elegans* (**Figure 2**). Chemosensory neurons that have been principally studied for roles in olfactory learning and navigation – AWA, AWB, AWC, and ASE – are the most broadly tuned neurons, with high sensitivity to many different types of molecules. AWA is comparatively more strongly activated by ketones, AWB by some esters, and ASE by alcohols. AWC is inhibited by every odorant that we tested. Other olfactory neurons – such as ASK, ASJ, or ASG – are more narrowly tuned, activated by a small number of structurally similar odorants (**Figures 4, S5**).

When the activity patterns of all broadly and narrowly tuned chemosensory neurons are taken together, a highly reproducible and distinct spatial map of neuronal activity emerges for each olfactory stimulus. This map encodes both odorant identity and intensity across the space spanned by our panel of 23 diverse chemicals tested at multiple concentrations (**Figure 5**).

How might *C. elegans* use an ensemble-level code for olfaction? Broadly tuned neurons permit coarse identification of odorants. Each narrowly tuned neuron is sensitive to a smaller region of olfactory space. When a narrowly tuned neuron is active, the possible identities of each olfactory stimulus are limited to those odorant molecules inside its region of sensitivity. When a neuron is inactive, molecules inside its region of sensitivity are ruled out. Combinatorial activity patterns among chemosensory neurons with different regions of sensitivity can provide enough information to pinpoint the identity and concentration of an odorant stimulus. When these ensemble-level patterns are highly reproducible, accurate discrimination can be performed with single stimulus presentations. Ensemble-level codes may also improve robustness, compensating for trial-to-trial variability in the responses of individual chemosensory neurons.

### Diverse sensory neuron tuning properties could be the result of multiple receptors

If an ensemble-level code is used to discriminate odorants, it is the unique response properties of each chemosensory neuron that allows each to contribute information to the spatial activity map that encodes olfactory stimuli. Removing any chemosensory neuron will lower the accuracy of stimulus classification based on ensemble activity (**Figure 6**). Because ensemble-level activity is largely independent of synaptic communication between neurons in *C. elegans* (**Figure 3B**), the tuning of each chemosensory neuron is, for the most part, a cell-intrinsic property.

In *C. elegans*, the tuning of each chemosensory neuron is shaped by the expression and properties of multiple receptors, not by the sensitivity of a single receptor as is typical in larger animals. ODR-10, highly expressed in AWA, remains the only characterized olfactory receptor for diacetyl (36,53). However, AWA also responds to many other odorants in a manner that is independent of ODR-10, direct evidence that AWA expresses multiple receptors (15–17,35). Moreover, other sensory neurons that do not express ODR-10 are activated by diacetyl at higher threshold concentrations.

We uncovered a diversity of odorant dose-response curves in *C. elegans* (**Figure 2E, S3F**). This diversity is likely explained by the expression of multiple receptors in each chemosensory neuron. Variable dose-response curves across chemosensory neurons may reflect the cumulative activities of different sets of receptors with different binding affinities for a given odorant. Moreover, each chemosensory neuron tends to be sensitive to structurally similar odorant molecules, suggesting correlations in the chemical binding affinities of the receptors expressed by each neuron (**Figures 4, S5**).

We lack a comprehensive characterization of the repertoire of functional receptors expressed in each type of chemosensory neuron. This makes it difficult to quantitatively extract the molecular parameters of receptor-ligand interactions from dose-response curves, as has been done in other animals (4,62,63). We note that tuning to olfactory stimuli (defined as the fraction of the odor panel which elicit significant responses) does not appear to be correlated with the number of GPCRs expressed (13). For example, ADL expresses the most GPCR genes of any chemosensory neuron, but is sensitive to only 3 odorants in our panel. ASH, ASK and ASJ express many GPCR genes, but only ASH is broadly tuned to our odorant panel. ASE, another broadly tuned neuron, expresses the smallest number of GPCR genes.

One explanation for the lack of correlation between the number of expressed GPCRs and the breadth of tuning is that we do not know how many GPCRs are engaged in olfaction. For example, ADL, narrowly tuned for odorants but broadly tuned for pheromones, may use many of its GPCRs as ascaroside receptors. Moreover, tuning is shaped both by the number of receptor types and the spectrum of receptor properties. Until more receptors are comprehensively characterized, we cannot relate chemosensory tuning properties to GPCR expression patterns.

### Pheromone detection engages the chemosensory ensemble in distinct ways

Comparing ensemble-level responses to volatile odorants and pheromones, we found that chemosensory neurons that are more narrowly tuned to volatile odorants are more broadly tuned to pheromones (**Figure 7**). We do not know if the activation of pheromone-sensing neurons by volatile odorants reflects cross-reactivity of pheromone receptors to small organic molecules, or whether these narrowly tuned neurons express receptors that are specific to each stimulus class. We also do not know if the activation of broadly tuned olfactory neurons by pheromones reflects cross-reactivity of olfactory receptors to large organic pheromone molecules. In any case, widespread ensemble-level activity across all chemosensory neurons in response to odorants and pheromones encodes substantial information that can be used to accurately identify any chemical stimulus.

### Comparisons with olfactory systems in larger animals

In larger animals, each sensory cell typically expresses a single receptor type. When domains of sensory neuron activity in larger animals are represented in a chemical structural space, such as the one that we used for *C. elegans*, response domains tend to be clustered. Olfactory neuron ensembles span the full range of chemical space by connecting the clustered response domains of different olfactory sensory neurons (1–6).

In *C. elegans*, each sensory neuron extends its sensitivity across a contiguous region of chemical space (**Figure 4**). This suggests that each neuron is tuned to shared molecular properties, as opposed to being faithful “labeled-line” detectors of a set of unique odorants. The broad tuning of many *C. elegans* sensory neurons is probably caused by the combined activities of different receptors. Each receptor may be tuned to a smaller region of chemical structural space. Connecting the regions of chemical space sensed by each receptor could produce the broad region of chemical space sensed by each neuron. The tendency for even the most broadly tuned neurons to be most strongly activated by certain chemical classes suggests correlations in the cell-specific expression of receptor molecules. Another consequence of the multi-receptor nature of *C. elegans* sensory neurons may be their exhibition of graded responses over a broad dynamic range of concentration. As additional receptor types with higher thresholds are recruited at higher concentrations of a given odorant, a sensory neuron gradually and cumulatively becomes more active.

### Discrepancies with previously reported chemosensory responses

We have characterized >900 neuron-stimulus pairings, including many previously undescribed responses. Where our measurements overlapped with previous studies, we found general agreement with previously reported neuronal responses (see **Results**). However, we also observed some discrepancies.

We did not observe previously reported OFF responses in AWC. This might be due to two factors. First, to map the tuning properties of chemosensory neurons, we used stimulus conditions that would minimize adaptation. We presented odorants in short 10s pulses with long intervening blank periods between presentations. Previously reported OFF responses in AWC had been observed with longer odor stimulus presentations (15,41). Second, some previously reported OFF responses were observed in one of the two asymmetric AWC neurons. Here, we did not separate the responses of ^ON^ and AWC^OFF^ neurons, and so any asymmetric AWC response would be lost in the population average.

We also did not recapitulate some previously recorded sensory neuron responses to ascarosides (26–29). This may be due to differences in the age and sex of tested animals. To be consistent with our own volatile odorant experiments, we recorded from young adult hermaphrodites. Different ascaroside responses in previous reports were observed in males and juvenile hermaphrodites.

### Limitations and future studies

Calcium imaging provides a coarse-grained measure of neuronal activity. Here, we primarily quantified peak calcium responses, omitting differences in dynamics, spiking, or asymmetric responses, all of which likely encode additional information. Thus, our estimates of the information encoded in ensemble-level activity represent conservative lower bounds.

Our analysis of synaptic transmission mutants suggests that synaptic transmission is not the primary driver of ensemble-level responses (**Figure 3**). However, synaptic connections and feedback may still shape the magnitude and dynamics of neuronal responses in important ways. For example, it has been suggested that feedback by neuropeptide signaling causes ASE to respond when benzaldehyde is detected by other sensory neurons (17). This and other forms of non-synaptic signaling may also contribute to coordinated activity among chemosensory neurons.

How is ensemble-level information transformed into behavior? Downstream from the chemosensory ensemble, interneuron networks resemble both a reflexive avoidance circuit (consisting of the command interneurons AVA, AVB, and AVD that primarily receive inputs from ASH) and a circuit for learning and navigation (consisting of the interneurons AIA, AIB, AIY, and AIZ that integrate the activity of the entire chemosensory ensemble) (**Figure 1A**) (10,30,31,33,37,40,64,65). ASH might be part of a “nociceptive labeled line” that maps the detection of noxious stimuli to rapid escape responses. However, the output of the entire chemosensory ensemble appears to be integrated and decoded by another more complex interneuron network. Large-scale multineuronal recording methods (21,66,67) that extend from the chemosensory neurons to downstream interneurons are needed to understand how ensemble-activity is mapped to decision-making circuits and behavioral responses.

### Perspectives

The extent to which any animal –*C. elegans*, insects, or vertebrates – exploits the collective activity of chemosensory neurons to decode olfactory inputs is poorly understood. On one hand, the “dimensionality” of the olfactory code is often presumed to be as large as the number of distinct chemosensory neurons that contribute to the code (68). If so, the ability to detect even small numbers of different molecules, each with specificity to different subsets of chemosensory neurons, can create the potential to discriminate astronomical numbers of olfactory stimuli (69). On the other hand, animals may trade a high-dimensional olfactory coding strategy for one that allows for rapid and efficient identification of odorants. This can be accomplished using a small number of the earliest responding (or primary) olfactory receptors and neurons, as seen in recent experiments with rodents that explore “primacy models” of the olfactory code (70).

The diversity of activity patterns available to the chemosensory ensemble allows one-to-one mapping to a much larger number of odorant stimuli than neurons. Does *C. elegans* use all the information encoded in the combinatorial possibilities of its chemosensory ensemble to increase the variety of internal olfactory representations and outward behaviors? Answering this question requires high-dimensional measurements that extend from olfactory perception, as in this study, to decision-making circuits and behaviors. Combining high-throughput odorant stimulation with brain-wide imaging and tracking in behaving animals is becoming possible with advances in microfluidics and imaging (50,71–74). While the combinatorial possibilities of the olfactory code are still large in *C. elegans*, its relatively small size makes it a useful system to explore the relevance of ensemble-level olfactory codes.

## Methods

### Worm maintenance

All *C. elegans* lines used in this project were grown at 22°C on nematode growth medium (NGM) plates seeded with the *E. coli* strain OP50. All animal lines were allowed to recover from starvation or freezing for at least two generations before being used in experiments. All animals used in experiments were young adults.

### Plasmids and crosses

To construct the ZM10104 imaging strain we created and then crossed two integrated lines, one expressing GCaMP6s and one expressing the wCherry landmark. The first of these lines, ADS700, was made by co-injecting *lin-15(n765)* animals with pJH4039 (*ift-20* GCaMP6s::3xNLS) and a *lin-15* rescuing plasmid. A stable transgenic line (hpEx3942) with consistent GCaMP expression in the chemosensory neurons was selected for integration, and transgenic animals were irradiated with UV light to integrate the transgenes into the genome. The resulting integrated line (aeaIs008) was backcrossed four times against N2 wild type. The second line, ADS701, was similarly made by co-injecting *lin-15(n765)* animals with pJH4040 (*gpc-1* wCherry) and a *lin-15* rescuing plasmid. A stable transgenic line with good wCherry expression was selected for integration, and transgenic animals were irradiated with UV light to integrate the transgenes into the genome. The resulting integrated line (hpIs728) was backcrossed four times against N2 wild type. To make ZM10104, ADS700 hermaphrodites were crossed with N2 males. Heterozygous aeaIs008/+ male progeny were then crossed with ADS701 hermaphrodites. F1 progeny were picked for wCherry expression, and F2 progeny were picked for both GCaMP6s and wCherry expression. The line was then homozygosed in the F3 generation.

The ADS707 mutant imaging line was created by crossing the ZM10104 line with EG9631, an *unc-13(s69)* mutant obtained from the CGC (55). EG9631 hermaphrodites were crossed with ZM10104 males. Heterozygous (aeaIs008/+; hpIs728/+; *+/unc-13)* F1 hermaphrodite progeny were selected by GCaMP6s and wCherry expression and wild type locomotion (*unc-13* is recessive). F2 progeny were picked for fluorescence and the *unc-13* uncoordinated phenotype. The line was homozygosed for fluorescence in the F3 generation.

### Microfluidics

We used a modified version of a microfluidic system capable of delivering multiple odors to *Drosophila* larvae (4). The microfluidics chip is designed with an arbor containing delivery points for multiple stimuli, together with a buffer delivery point and two control switches, one for buffer and one for odor (**Figure 1B**). At any given time, three flows are active: one of the control switches, the buffer blank, and one odor stimulus. The chip is designed to maintain laminar flow of each fluid, and the flow is split between a waste channel and an odor channel which flows past the animal’s nose. The chip described here is designed to switch rapidly from one stimulus to the buffer. After the flows pass the animal, they exit the chip via a waste port at atmospheric pressure. Waste is removed with a vacuum.

We grafted the odorant delivery arbor to a *C. elegans* loading chamber similar to those designed by Chronis, et al. (50). We designed a loading chamber suitable for adult *C. elegans*, a narrow channel 62 μm wide and 30 μm high, with a gently tapered end. The tapered end serves as a guide to help hold the animal’s nose in place without distorting the animal. The microfluidic device pattern was designed in AutoCAD, and the design was translated to silicon wafer using photolithography. The photomasks of the design were printed using CAD/Art Services, Inc. The silicon wafer was then used as a mold for polydimethylsiloxane (PDMS) to fabricate microfluidic devices. The PDMS components were then removed from the silicon wafer, cut to size, and had access channels made with a biopsy punch. The completed PDMS components were then plasma bonded to No. 1 glass cover slips. To minimize contamination from dust, all microfluidics assembly was done in a cleanroom.

### Preparation of odorant and buffer solutions

Odorants were diluted in CTX buffer (5 mM KH_2_PO_4_/K_2_HPO_4_ at pH 6, 1 mM CaCl_2_, 1 mM MgSO_4_, 50 mM NaCl, adjusted to 350 mOsm/L with sorbitol). To prevent contamination, each odor condition was mixed and stored in its own glass bottle, and delivered through its own glass syringe and tubing. Furthermore, a new microfluidic device was used for a single consistent panel of odors. The single ascarosides (25) were diluted in CTX buffer to 200 mM concentration for presentation to the animals.

### Imaging setup

We used a single-photon, spinning-disk confocal microscope to capture fluorescent images from intact *C. elegans*. The microscope was inverted to allow for easy access to the microfluidics device mounted on the stage. We employed a 488 nm laser to excite GCaMP *in vivo*, and used a 561 nm laser to excite the wCherry landmark. To minimize cross-talk between channels, lasers were fired sequentially during multicolor recordings. We captured images with a 60× water-immersion objective with an NA of 1.2. Volumes were acquired using unidirectional scans of a piezo objective scanner. All fluorescence microscopy is a trade-off between spatial resolution, temporal resolution, laser power, and signal strength. We optimized two sets of imaging conditions, one set for activity imaging and another set for landmark imaging. Both sets of imaging conditions capture the region containing the majority of the neurons in the head of *C. elegans*, a volume of 112 μm by 56 μm by 30 μm.

In any given experiment, acquisition of a landmark volume precedes acquisition of an activity movie. This volume, which contains both green and red channels, allows us to identify neurons of interest. The spatial resolution of these volumes is 0.5 μm × 0.5 μm × 1.5 μm/voxel, with the z-resolution of 1.5 μm set by the point spread function.

The activity movies were acquired at a high speed in the green channel only, with lower spatial resolution (1 μm × 1 μm × 1.5 μm/voxel). At this resolution, we could acquire volumes at 2.5 Hz in standard acquisition mode.

### Analyzing multi-neuronal recordings

The neurons in each activity recording were identified and then tracked through time using a neighborhood correlation tracking method. The criteria for identifying each neuron class are described in Appendix A. Neurons which could not be unambiguously identified were excluded from the dataset. All neuron tracks were then manually proofread to exclude mis-tracked neurons. Activity traces were bleach corrected and reported in:

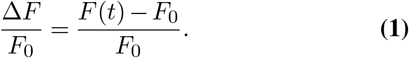

Normalization by baseline fluorescence *F*_0_ allowed for direct comparisons within a given neuron class across L/R and across individuals. The baseline *F*_0_ value was determined individually for every recorded neuron, set at the 5th percentile of the distribution of bleach-corrected fluorescence values, with the opportunity for manual correction.

We employed 2-tailed, paired t-tests to compare the mean signal during stimulus presentation with an unstimulated period of identical length within the same neuron. Neurons were tested for both ON and OFF responses. The p-values were corrected for multiple testing using FDR (75). To test for asymmetric neuron responses, we used 2-tailed, two-sample t-tests (unpaired).

## AUTHOR CONTRIBUTIONS

A.L., S.Q., H.C., V.V., M.Z. and A.S contributed to writing this manuscript. A.L., V.V., and A.S. designed the experiments. A.L., M.W., W.H., and M.Z. designed and built the transgenic C. *elegans* strains used in this project. A.L. and V.V. designed and built the imaging setup, microfluidics devices, and the wrote the software to extract neural traces. A.L., H.C., G.C., N.T., and R.V. carried out the imaging experiments and analyzed the data. A.L., S.Q., and C.P. developed the theoretical models. L.L. prepared some of the pheromone reagents.

## ACKNOWLEDGEMENTS

We thank Guangwei Si and Jessleen Kanwal for their advice on microfluidics design and operation and Maedeh Seyedolmohadesin and Mahdi Torkashvand for our discussions on ascaroside responses. We also thank Sandeep Robert Datta, Steven Flavell, and the members of the Zhen, Pehlevan, and Samuel labs for their advice on the project and comments on the manuscript. The EG9631 strain was obtained from the CGC, which is funded by the NIH Office of Research Infrastructure Programs (P40 OD010440). Microfluidics devices were manufactured using the Soft Materials Cleanroom facility of the Harvard MRSEC (DMR-1420570). This work was supported by NSF Brain Eager (NSF IOS-1452593), NSF Physics of Living Systems (NSF 1806818), NSF Ideas (NSF IOS-1555914) and the Canadian Institute of Health (CIHR Foundation Scheme 154274) grants.

## Supplement Supplemental methods

## A: Identifying neurons in the ZM10104 strain

The ZM10104 strain used in this experiment expresses two fluorescent proteins: GCaMP6s driven by the *ift-20* promoter, and wCherry driven by *gpc-1*. GCaMP6s expression was localized to neuronal nuclei to minimize spatial overlap of neighboring neurons, and to make identification of neurons easier. The promoter *ift-20* drives GCaMP expression in all ciliated sensory neurons. Our neurons of interest, the chemosensory neurons, lie in the lateral ganglia, but note that this promoter is pan-sensory, driving expression in cells outside of the lateral ganglia. The wCherry landmark is expressed in the cytoplasm of AFD, AWB, ASI, ASE, AWC, and ASJ. Note that it also is expressed in RIB, a neuron which is not labeled with GCaMP.

Relative positions are given in the orientation in **Figure S1**, with the nose to the left, the tail to the right, dorsal top, and ventral bottom. Relative positions should be interpreted as “usually but not always,” unless otherwise noted. Also note that overly compressing an animal in any direction will distort the relative positions. Before identifying neurons, it is important to identify the orientation of the animal in the recording by figuring out where the dorsal-ventral (DV) plane lies. This is most easily done by identifying the plane of bilateral symmetry. Once you have oriented yourself, you can begin to identify neurons.

The easiest neurons to immediately identify in this strain are ASH, ASJ, and the anterior “triplet” of ASK, ADL, ASI. It is often effective to identify these neurons first, then work on the other neurons using the color landmarks and process of elimination. AWC and ASE should usually be in the neighborhood of ASH, and the four neurons AWA, AWB, ADF, and ASG are between ASH and the anterior triplet. These four neurons occasionally overlap. To avoid signal mixing, overlapping neurons were excluded from the dataset. For each odorant condition, neuronal identification was carried out independently by at least two individuals.

**Figure S1.**
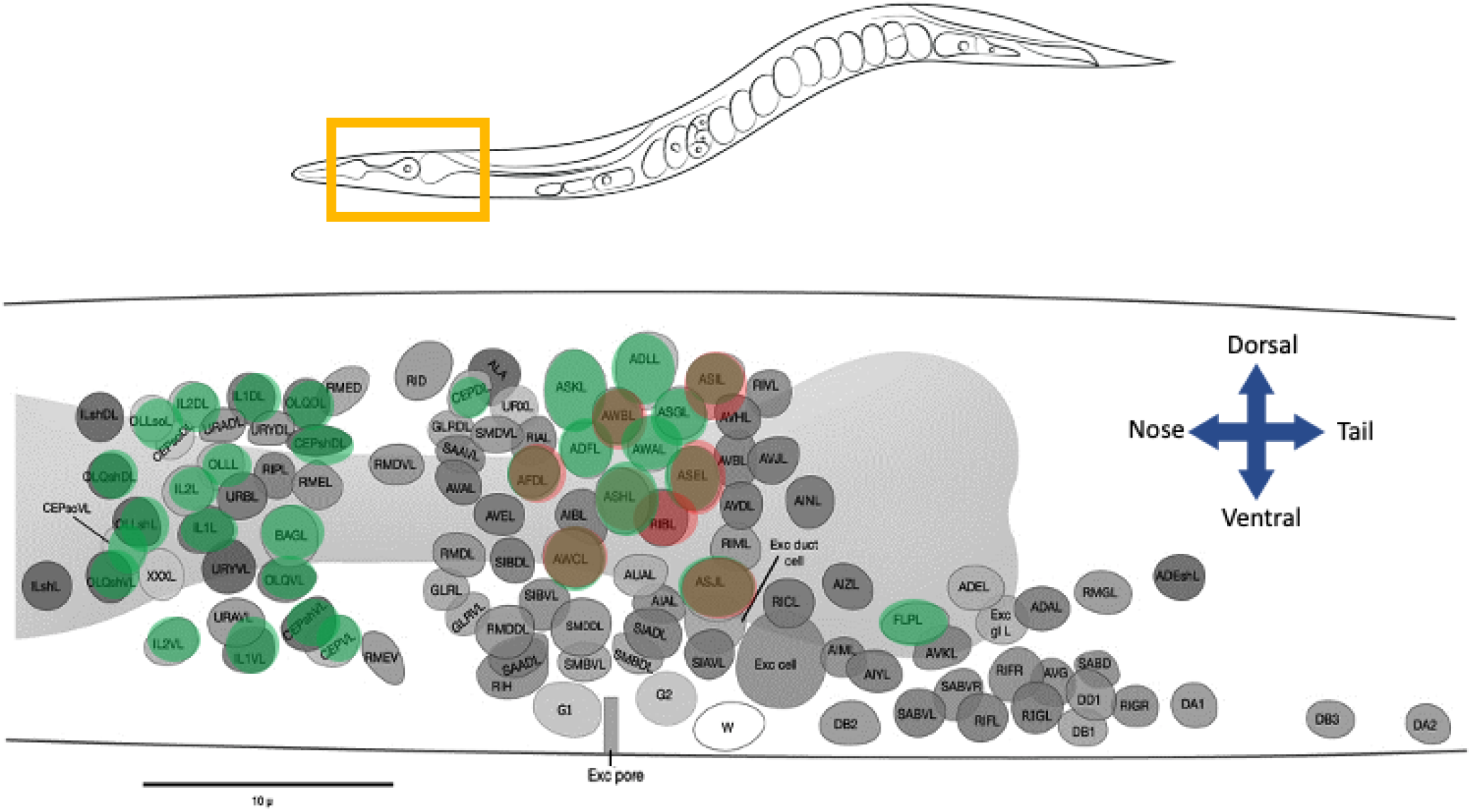
Identifying neurons in the ZM10104 strain. The *ift-20* promoter drives GCaMP expression in the nuclei of ciliated sensory neurons. The nuclei of the chemosensory neurons are all posterior to the nerve ring. A red landmark is provided by cytoplasmic expression of wCherry in the neurons AFD, AWB, ASI, ASE, AWC, and ASJ. Underlying *C. elegans* figure adapted from the digital version of White et al. 1986 (Wormbook)(48).

### Criteria for identifying each neuron class

| Neuron | Color(s) | Relative Position | Notes |
| --- | --- | --- | --- |
| ASK | green | leftmost of the anterior triplet | large. do not confuse with URX, a small oblong neuron above ASK |
| ADL | green | part of the anterior triplet | larger than ASI |
| ASI | green & red | part of the anterior triplet | use color to distinguish from ADL |
| ASH | green & red | left of ASE, below AWA | bright, circular |
| ASE | green & red | right of ASH | smaller than ASH |
| AWC | green & red | variable. below ASH but can be to the left, directly below, or to the right | often oblong in shape |
| ASJ | green & red | tail end of the ganglion, bottom left | distance from AWC can vary |
| AWA | green | variable. usually above ASH | smaller than ASH, circular |
| AWB | green & red | position variable, usually directly below the anterior triplet | small, dim, a bit oblong. use color to identify |
| ADF | green | usually left of AWA, AWB | dim |
| ASG | green | usually right of AWA, AWB | small, circular |

To minimize the chances of incorrect identification, neuronal IDs for each odorant condition were reviewed by at least two individuals, and ambiguous neurons were omitted from the analyzed datasets.

## B: List of odorants

| Odorant | Chemical class | Behavioral valence (low conc.) |
| --- | --- | --- |
| 1-pentanol | alcohol | attractive |
| 1-hexanol | alcohol | attractive |
| 1-heptanol | alcohol | repulsive |
| 1-octanol | alcohol | repulsive |
| 1-nonanol | alcohol | repulsive |
| isoamyl alcohol | alcohol | attractive |
| geraniol | alcohol | attractive |
| benzaldehyde | aromatic | attractive |
| methyl salicylate | aromatic | attractive |
| ethyl acetate | ester | attractive |
| ethyl butyrate | ester | attractive |
| ethyl heptanoate | ester | attractive |
| pentyl acetate | ester | attractive |
| butyl butyrate | ester | attractive |
| diacetyl | ketone | attractive |
| 2-butanone | ketone | attractive |
| 2-heptanone | ketone | attractive |
| 2-nonanone | ketone | repulsive |
| 2,3-pentanedione | ketone | attractive |
| 2,5-dimethylpyrazine | pyrazine | attractive |
| 2-methylpyrazine | pyrazine | attractive |
| 2-isobutylthiazole | thiazole | attractive |
| 2,4,5-trimethylthiazole | thiazole | attractive |

## C: Neuron tracking and signal extraction

To segment the neuronal nuclei in each recording, we built a GUI which allows users to navigate each 3D landmark image and click to add or remove neuron centers (21,73). This GUI allows the user to toggle between multiple fluorescent channels and a maximum projection, allowing the user to take advantage of any fluorescent landmark labels in the strain. Complete labeling of all neuron centers is only necessary once for a given animal, even if multiple recordings have been made. The user then labels a small handful of widely spaced neurons (4−8) in the first frame of the activity recording. This small number of labeled neurons helps the tracking algorithm to compensate for any global motion or distortion that may have occurred in the animal between the landmark volume and the activity movie. In addition to segmentation, the GUI allows neurons to be manually identified. The names the user applies are then associated with the activity traces of those neurons.

### Neighborhood correlation tracking of individual neurons

While the entire brain of the worm can distort significantly across large distances, the neighborhood immediately surrounding a neuronal nucleus of interest tends to remain consistent, with little local deformation. Our image registration strategy relies on this fact. Instead of attempting to identify neuron centers in every frame, we try to match the neighborhood surrounding the neuron center in the first frame to the most similar neighborhood in the following frame. We then return the center of the new neighborhood as the position of the neuron center in the next frame.

We first employ this approach to map the neuron centers identified in the high-resolution landmark volume during the segmentation step onto the first frame of the activity movie, which is captured at a lower resolution. We then proceed to compare each frame of the movie to the next. The neighborhood correlation comparison is made independently for each neuron. While we lose some information about local deformations by not integrating information about how neighboring neurons are moving, we gain the ability to run the tracking of each neuron in a dataset as a parallel process, dramatically decreasing runtime. This also prevents a mistake in tracking one neuron from propagating to other nearby neurons. We run the tracking on a down-sampled version of the activity movie, also to improve runtime.

For a given neuron center, the tracking algorithm goes through the following steps:

1. Given the position of the given neuron center in the current frame, *n_t_* = (*x_t_, y_t_,z_t_*), we identify the neuron’s local 3D neighborhood *N_t_* in that frame, the volume with dimensions 2*a* * 2*b* * 2*c*, in the region spanned by [*x_t_* – *a*, *x_t_* + *a*], [*y_t_* – *b, y_t_* + *b*], and [*z_t_* – *c,z_t_* + *c*].
2. We identify the naive center in frame *t* + 1, from where we begin our search for the neighborhood most similar to *N_t_*. For the first frame of the movie, this point is adjusted by a distance-weighted average of the manually labeled neurons: 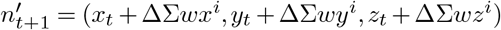. For any other frame, we simply take the naive center as the center of the previous frame, 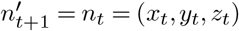.
3. Starting from the naive center 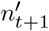, we perform image registration between the maximum intensity projections in *x*, *y*, and *z* of putative neighborhood 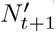 and the previous neighborhood *N_t_*, computing the pairwise correlation of these images. We then repeat this process, moving the putative center 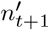 by 1 pixel per iteration until one of the following occurs:

a. The algorithm finds a putative neighborhood 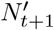 whose correlation with *N_t_* exceeds the confidence threshold *C* (usually set at above 90%). This putative neighborhood is then defined as *N_t_*_+1_.
b. The algorithm tests all putative neighborhoods within a maximum search radius *r*_max_ of the naive center 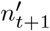, but failed to find a putative neighborhood whose correlation exceeds the confidence threshold *C*. The algorithm then returns the putative neighborhood with the highest correlation with *N_t_* as *N*_*t*+1_.
c. If no neighborhood is found with a correlation exceeding a minimum value, the neuron is considered lost in frame *t*+ 1, likely either due to motion taking the neuron outside the region of interest. No center is reported, and the last reported neighborhood *N_t_* is used as the basis of comparison for following frames (*t* + 2, *t* + 3, etc.).
4. The center of neighborhood *N*_*t*+1_ is defined as the neuron center in this frame, *n*_*t*+1_.
5. Repeat until the end of the activity movie is reached.

We can optimize the tracking parameters such as neighborhood size (*a, b, c*), maximum search radius *r*_max_, and confidence threshold *C* for both accuracy and speed for different imaging conditions.

### Extracting calcium dynamics

To extract calcium signals, we first map the positions of each tracked neuron center back onto the original-resolution volumetric images. We then extract fluorescence values from these images. We identify a small volume around each neuron center, containing voxels whose fluorescence will be assigned to the neuron. This volume is set as 2 μm × 2 μm × 3 μm for our data. We compute the mean of the 10 brightest pixels within this volume to extract a raw fluorescence trace *F_r_* (*t*). We then account for photobleaching by exponential detrending, giving us a clean fluorescence activity trace *F*(*t*). We then identify the background fluorescence *F*_0_ for each neuron, and report normalized neuron activity Δ*F*/*F*_0_.

### Manual proofreading of traces

Manual proofreading is an opportunity to improve data quality by removing neurons which have been mistracked, adjusting the computer-determined baseline fluorescence *F*_0_, and correcting or adding nuclear IDs. Proofreading also enabled us to remove traces which were contaminated by signals from neighboring neurons. The software then compiles all processed traces for a given individual into a single data structure.

## D: Imputing missing single-trial responses

Across trials of all neurons and all conditions, about 20% of the neuron responses were either not captured, or excluded due to tracking mistakes or signal contamination issues. To perform single-trial discrimination analysis in the (*N*-dimensional) neural response space, we first had to fill these missing data points in a reasonable and biologically motivated way.

For a given odorant and *M* trials, the peak responses of the *N* = 11 sensory neurons can be compiled in a matrix 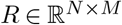. Without any assumptions for the values *R*, it is impossible to infer the missing data. Fortunately, due to the intrinsic correlation between the responses of different olfactory neurons, the full response matrix *R* is low rank (as indicated by the PCA of neural responses). We can use this low-rank information to recover the missing entries: “matrix completion” algorithms can solve this problem very efficiently (76,77).

To verify that matrix completion can indeed recover the missing entries faithfully, we performed a holdout evaluation. For the response matrix to each odor, we performed matrix completion after randomly removing 20 entries (*x_i_*, *i* =1, ⋯, 20). The imputed matrix is denoted as *X*\*. We then calculated the Pearson correlation coefficient *ρ* between the estimated entries 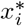 with the true entries *x_i_*. The average value of *ρ* is around 0.7 (**Figure**). We used the MATLAB code provided in (78) with default parameters for matrix completion (https://github.com/udellgroup/Codes-of-FGSR-for-effecient-low-rank-matrix-recovery). Specifically, we chose an algorithm based on minimization of the nuclear norm `MC_Nulcear_IALM`.

## E: Computational methods for discriminability quantification

For binary classification of all odorant pairs, we used linear regression and a simple SVM (linear or Gaussian kernel). To decode odor identity from the entire single-trial dataset, we built a multi-class classifier. We concatenate all of the single-trial responses of the 23 odorants at high concentration. Each trial is an 11-dimensional point, one dimension for every neuron class. Each point has an associated label indicating the odorant identity. This data set was randomly divided into 10 parts, 9 of which are used as a training set (90%) and one which is used as a validation set (10%).

We used the MATLAB function `fitcecoc` to fit a multi-class model which supports both SVM and other classifiers. Mechanistically, this method reduces the problem of overall classification into a sequence of binary classification problems. The performance was quantified by the classification error, estimated using the `crossval` function. The confusion matrix was generated using the functions `kfoldPredict` and `confusionchart`. The training is repeated 10 times, using each of the 10 parts of the datasets as the validation set, and the results were compiled.

For the *in silico* knockouts, we removed neurons from the training dataset, resulting, for example in 10-dimensional responses when one neuron was removed. We trained the multi-class classifier as above.

## F: Statistics, code, and software

All statistical computations and image analysis code were written and run in MATLAB using standard toolboxes, with the exception of the OME Bio-Formats API (used to read Nikon ND2 file formats) (79) and CET Perceptually Uniform Color Maps (80).

## Supplemental figures

**Figure S2.**
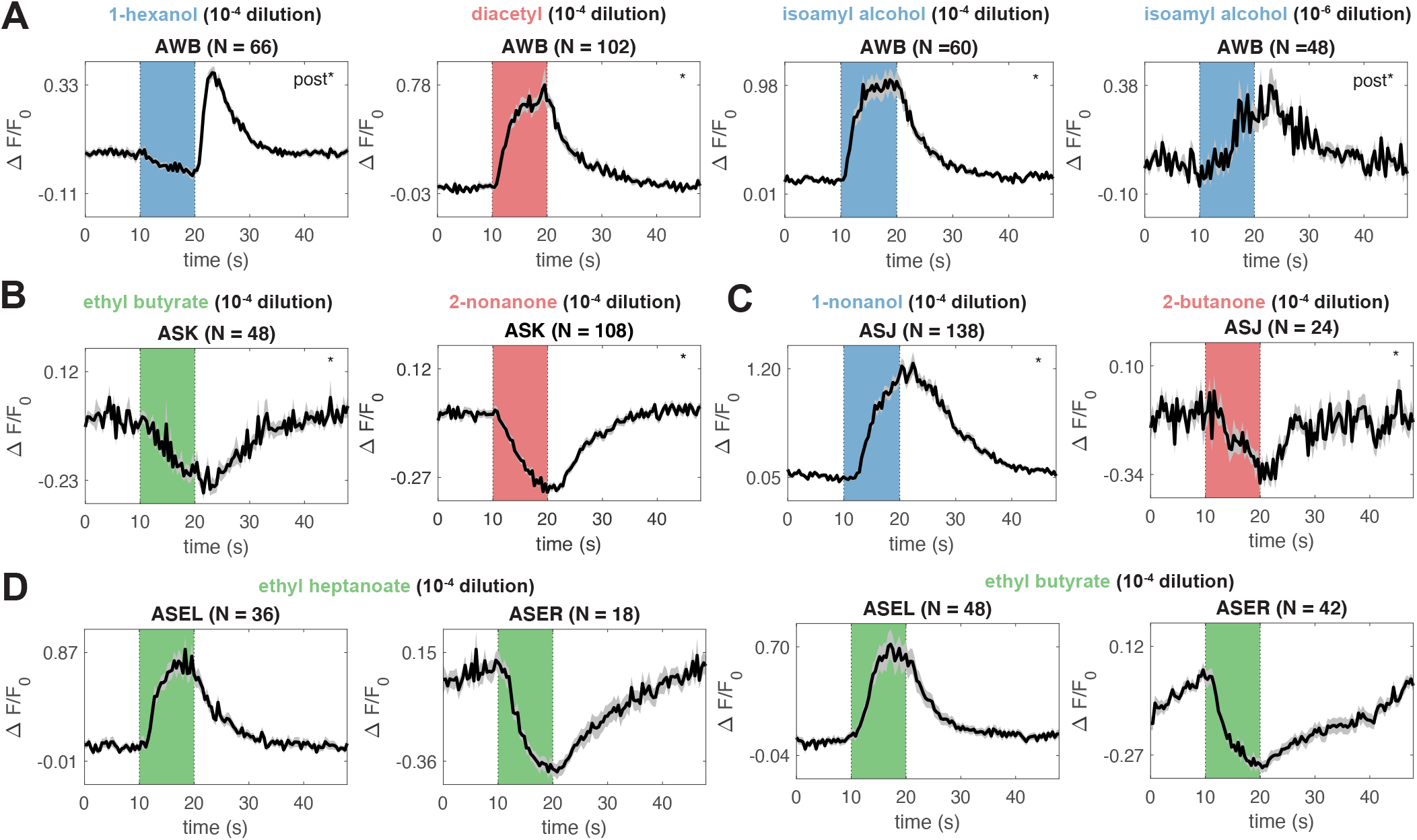
Single neuron response observations. **(A)** AWB is an OFF response for most stimuli, such as 1-hexanol, but is occasionally an ON response, as is the case for high concentration diacetyl. High concentration isoamyl alcohol elicits an ON response from AWB, but low concentration isoamyl alcohol elicits an OFF response. This has been previously observed in Yoshida et al., 2012 (15). **(B)** We observe inhibitory responses to some odorants in ASK. **(C)** ASJ has an excitatory response to some odorants, such as 1-nonanol, but has an inhibitory response to 2-butanone. **(D)** We observe L/R asymmetries in ASE in response to several odorants, such as ethyl heptanoate and butyl butyrate.

**Figure S3.**
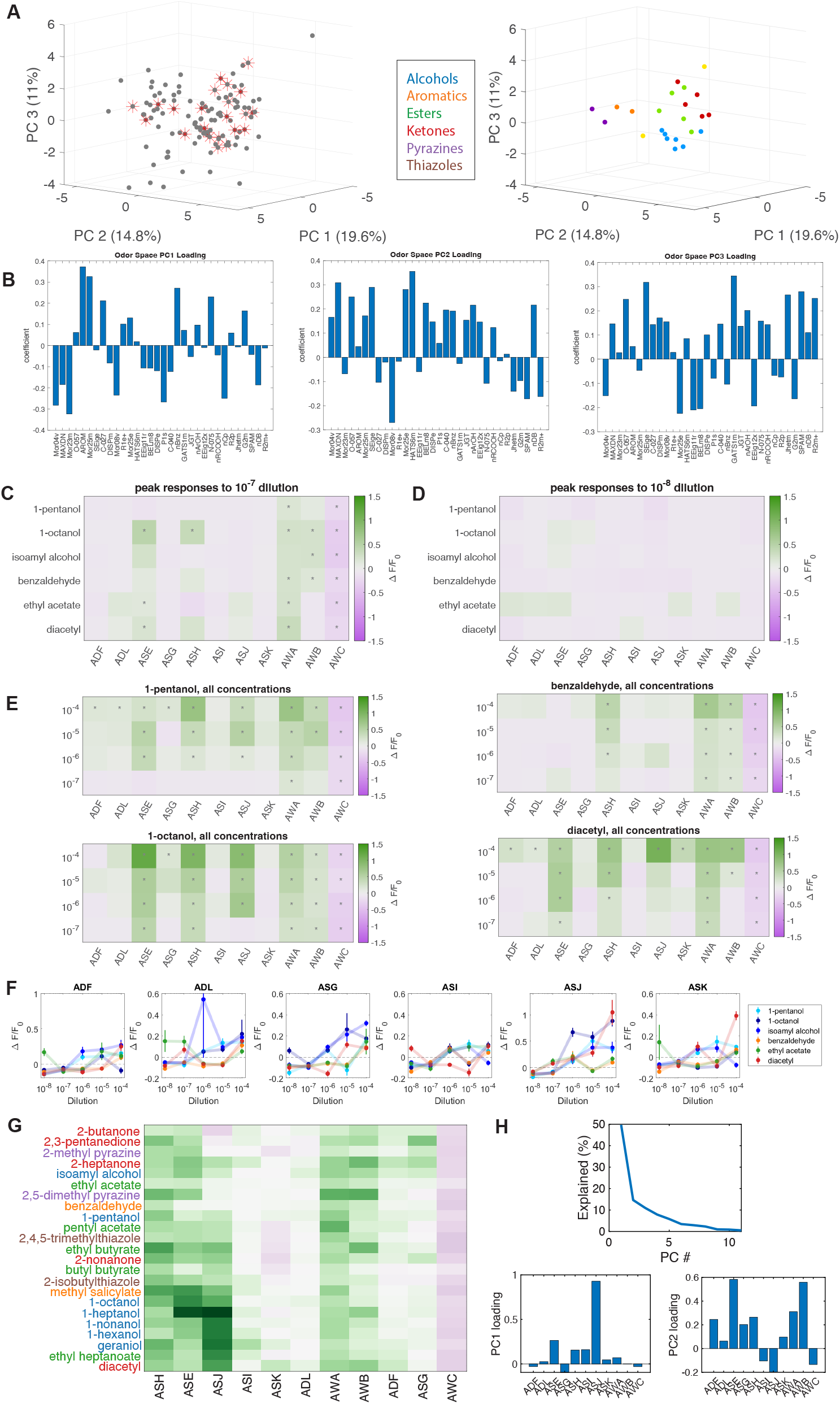
Supplemental panels for Figure 2. **(A)** An odor space constructed from the molecular descriptors of 122 odorants (gray) previously studied in *C. elegans*. We selected for our experiments a panel of 23 odorants (red) which span the odor space (left). On the right, these 23 odorants are presented in odor space colored by their chemical class. **(B)** The molecular descriptor loadings of the first 3 principal components of the *C. elegans* odor space, plotted on the same axes. The leading components of PC 1 are measures of aromaticity, and the leading components of PC2 are measures of electronegativity. Peak responses for six odors tested at **(C)**10^−7^ and **(D)** 10^−8^ dilutions. Statistically significant responses (*q* ≤ 0.01) are indicated with stars—no significant activity was observed at the lowest tested dilution. **(E)** Compiled responses to three representative odorants at multiple concentrations (1-pentanol, 1-nonanol, and benzaldehyde) show similar neural responses across concentration. The magnitude of neuron responses generally increases with increasing concentration, and for some conditions, additional neurons are recruited at high concentration. **(F)** Dose responses for the six sensory neurons not printed in **Figure 2E**. **(G)** Odorants (high concentration) clustered by their peak average neuronal responses. **(H)** The variance explained and the loadings of the first two principal components of the standardized average peak neural response PC space in **Figure 2F**.

**Figure S4.**
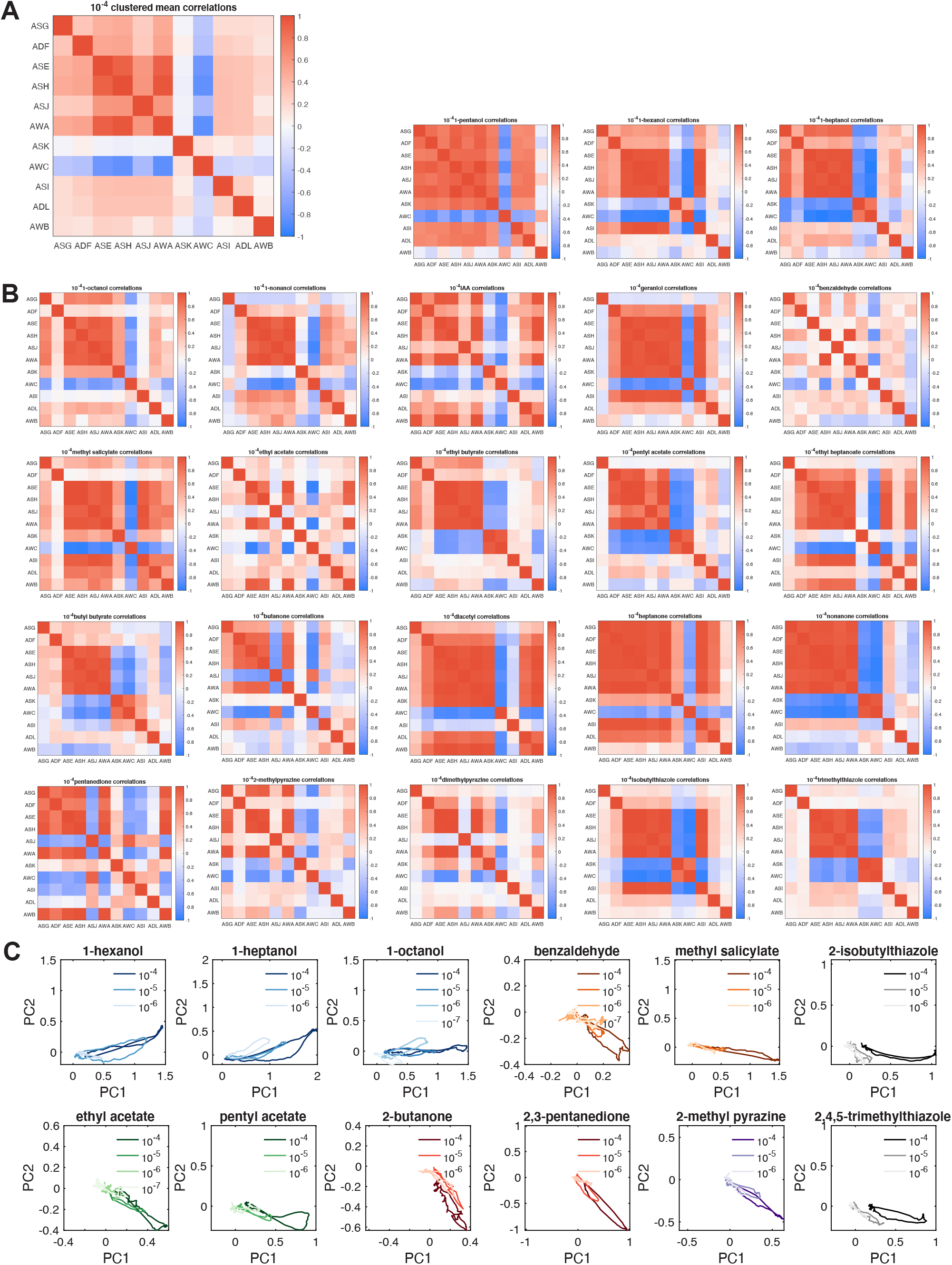
Time trace correlations and phase trajectory analyses. **(A)** Average time trace correlation map of the 11 chemosensory neuron responses across all 23 odorants. **(B)** Average correlation maps of responses to all 23 odorants at high concentration, plotted on the same axes, show diverse response dynamics. **(C)** Phase trajectory plots of average neural activity for select odorants, all plotted in a common PC space. The shade of each color indicates concentration, with low concentration indicated by a light shade and high concentration indicated by a dark shade. Different concentrations of the same odorant tend to generate similar trajectories.

**Figure S5.**
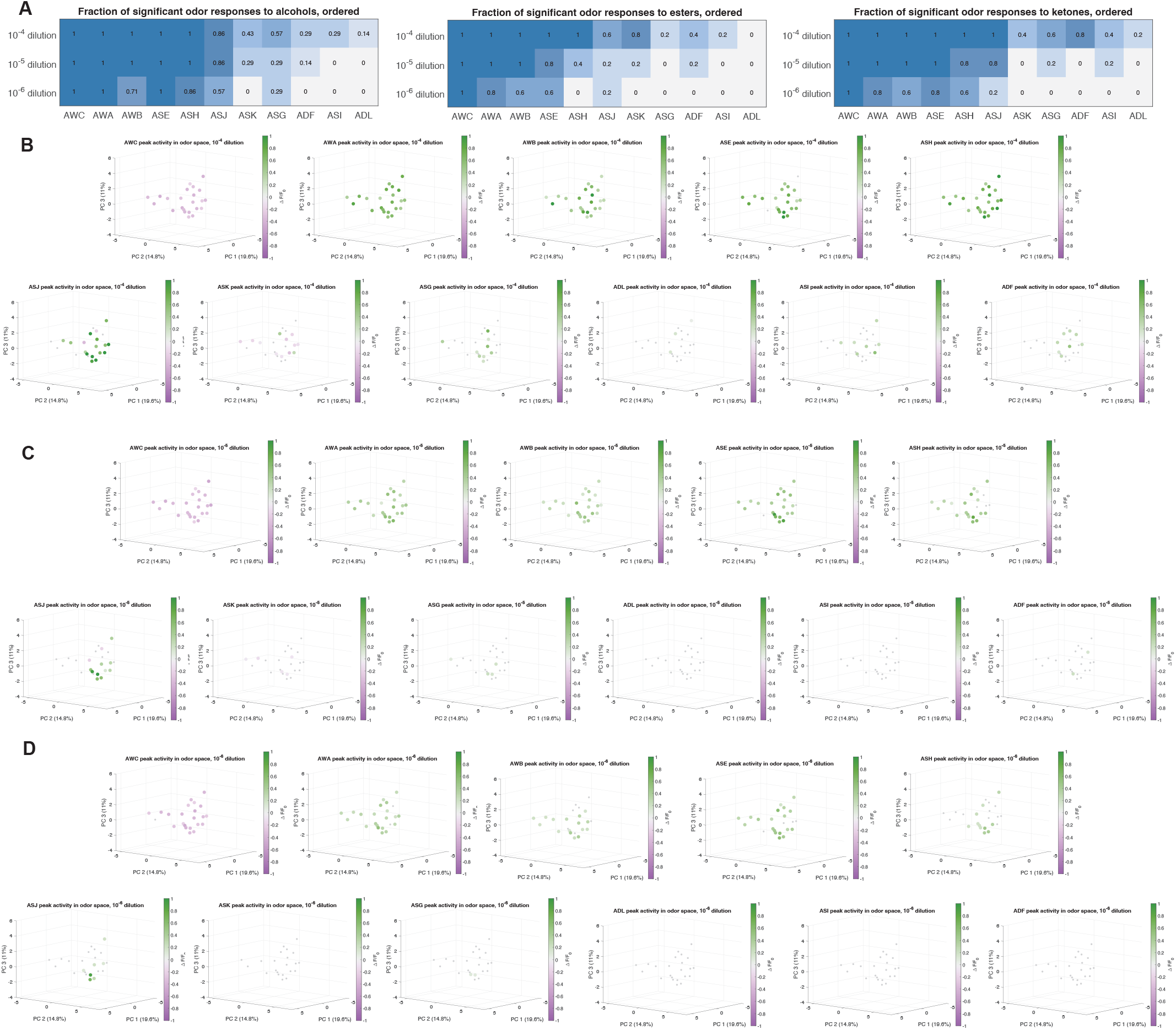
Average peak responses plotted in odor space. **(A)** The fraction of significant odor responses to three chemical groups: alcohols (7 total stimuli), esters (5 total stimuli), and ketones (5 total stimuli). Average peak responses of each of the 11 chemosensory neuron classes plotted in odor space (**Figure S3A**), at **(B)** high odorant concentration (10^−4^), **(C)**medium odorant concentration (10^−5^), and **(D)**low odorant concentration (10^−6^).

**Figure S6.**
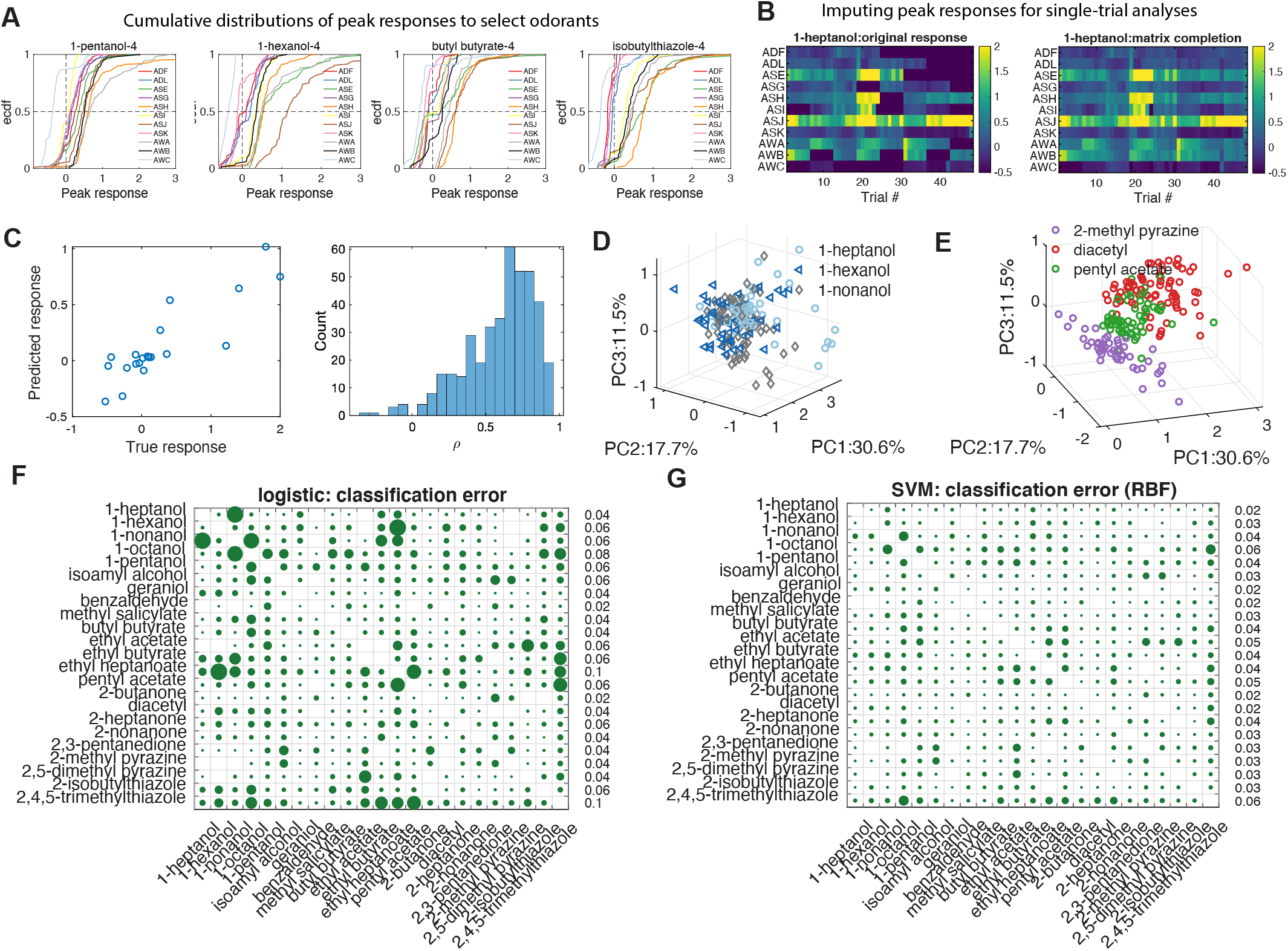
Supplemental panels for Figure 5. **(A)** Cumulative distributions of peak responses of every neuron (four exemplar odorants shown). **(B)** Signals were not always captured from all 22 chemosensory neurons in every trial. We used a matrix completion algorithm to impute these missing data points. Here are shown the peak responses all chemosensory neurons to 1-heptanol in different trials, with missing responses in black (left) and after matrix completion (right). **(C)** *Left*: To quantify the performance matrix completion, we randomly removed 20 measured responses (true response) and compared the imputed values from matrix completion (predicted responses).Right: The histogram of Pearson’s correlation coefficient between true responses and predicted responses. For each response matrix, we repeated 5 times. **(D/E)** Representations of single-trial peak neural responses to sets of **(D)** three similar and **(E)** three dissimilar odorants. These data are plotted in a PC space constructed from the individual trial responses to all odorants in the dataset. **(D)** We see that three similar odorants, the straight-chain alcohols 1-hexanol, 1-heptanol, and 1-nonanol, have more similar neural representations. **(E)** In contrast, three odorants of three distinct chemical classes, 2-methylpyrazine (a pyrazine), diacetyl (a ketone), and pentyl acetate (an ester), have more easily separable neural representations. Binary classification of all odorant pairs by **(F)** logistic regression and **(G)** SVM. Both methods return very low classification errors, demonstrating that the single-trial peak responses of any two odorants are linearly separable. Shown here are classification error heatmaps at high concentration (10^−4^ dilution), for which the average classification error is 0.055 for the logistic regression and 0.035 for the SVM.

